# A symbiotic MLO gene regulates root development via RALF34-triggered Ca^2+^ signalling in *Lotus japonicus*

**DOI:** 10.1101/2025.09.18.676995

**Authors:** Filippo Binci, Giacomo Guarneri, Sofía Cristina Somoza, Filippo Vascon, Arianna Capparotto, Edoardo Di Nuzzo, Giacomo Rago, Barbara Baldan, Laura Cendron, Lorella Navazio, Marco Giovannetti

## Abstract

*Mildew Locus O* (*MLO*) genes, initially identified as powdery mildew susceptibility factors, are increasingly recognized as multifunctional regulators implicated in diverse processes, including plant reproduction, root thigmotropism, and interactions with beneficial microbes. Recent evidence shows that MLO proteins can act as Ca^2+^-permeable channels in response to Rapid Alkalinization Factors (RALF) peptides in reproductive cells, pointing to broader roles in Ca^2+^-mediated signalling.

In this study, we investigate the symbiotic clade IV member LjMLO4 in the model legume *Lotus japonicus*, focusing on its role in root development and responsiveness to LjRALF34 peptides. We show that *LjMLO4* expression is strongly induced in root cells colonized by arbuscular mycorrhizal (AM) fungi, yet loss-of-function mutants exhibit only subtle AM-associated phenotypes. Instead, we uncover a previously uncharacterized function of LjMLO4 as a regulator of primary root growth and lateral root formation, acting even in the absence of AM fungal colonization and in a Ca^2+^-dependent manner. Heterologous expression in *E. coli* confirms that LjMLO4 facilitates Ca^2+^ transport, while genetic and physiological assays demonstrate its contribution to LjRALF34-triggered root growth responses and Ca^2+^ signalling.

Together, these findings identify LjMLO4 as a molecular hub between peptide signalling, Ca^2+^ transport and root system architecture, highlighting how MLO proteins integrate developmental, nutritional and symbiotic cues.

## Introduction

*Mildew resistance gene Locus O* (*MLO*) was first identified in barley as the genomic region where loss-of-function mutations confer resistance to *Blumeria graminis*, the causal agent of powdery mildew (Büschges et al. 1997). Following the cloning of the first *MLO* gene, phylogenetic analyses revealed that the *MLO* gene family is broadly conserved across the green lineage and diversified during evolution into seven distinct sub-clades (Devoto et al. 2003). Over the past two decades, members of this family have been implicated in diverse developmental and physiological processes in plants, such as plant reproduction (clade III), root thigmomorphogenesis and root hair elongation (clade I and II), interaction with biotrophic and necrotrophic pathogens (clade IV and V) as well as association with endophytes and arbuscular mycorrhizal fungi (clade IV) (Acevedo-Garcia et al. 2014; Jacott et al. 2021; Ogawa et al. 2025; Li and Xiao 2026). Despite this functional diversity, MLO proteins share conserved structural features: seven transmembrane domains, four cysteines in the extracellular/luminal space, a calmodulin (CaM)-binding domain and a crucial tryptophan residue at the cytosolic carboxy-terminus. Moreover, a few conserved motifs suggest that MLOs may be involved in transmembrane transport (Kusch et al. 2016).

At the cellular level, MLOs have been associated with processes such as programmed cell death, callose deposition, vesicular trafficking and modifications of cell wall composition (Acevedo-Garcia et al. 2014; Kusch et al. 2019; Hübbers et al. 2023). A unifying feature across clades is their functional dependence on Ca^2+^, as suggested by the presence of the CaM-binding domain and the need for external Ca^2+^ to fulfil their functions (Kim et al. 2002; Chen et al. 2009; Meng et al. 2020; Jacott et al. 2021). Recently, the long-sought biochemical activity of MLOs has been unravelled via patch-clamp assays, showing that MLOs are a novel plant-specific family of cation-permeable channels, with a preference for divalent cations such as Ca^2+^ and Mg^2+^, and that they are activated by Rapid Alkalinization Factors (RALF) peptides (Gao et al. 2022, 2023). For example, in *Arabidopsis thaliana* synergid cells, clade III MLO7/NORTIA perceives pollen tube-derived RALF4/RALF19 through interaction with the CrRLK1L-type receptor-like kinase FERONIA and the glycosylphosphatidylinositol-anchored protein LORELEI, thereby becoming an active cation-permeable channel (Gao et al. 2022). The functional interaction between FERONIA and MLOs was proposed before (Kessler et al. 2010) and ectopic expression of a constitutively active MLO (faNTA) has recently been found to restore Ca^2+^ oscillations in root hairs of *feronia* mutants (Ogawa et al. 2025). These findings open new avenues of research in the understanding of the cellular function of MLO as a Ca^2+^-permeable channel downstream to the RALF/CrRLK1L-type signalling module. In parallel, Arabidopsis clade I MLO4 is required for the cytosolic Ca^2+^ elevation needed for re-directioning the root tip growth upon physical stimulation (Zhang et al. 2022).

Evidence from heterologous expression further indicates functional conservation across clades: barley clade IV MLO mediates Ca^2+^ transport in oocytes (Gao et al. 2022). Although clade IV *mlo* barley mutant is mainly known for resistance to powdery mildew, it also displays altered AM fungal (AMF) colonization (Ruiz-Lozano et al. 1999; Hilbert et al. 2020; Jacott et al. 2020). AM symbiosis is an ancient and widespread mutualistic interaction in which fungi of the subphylum Glomeromycotina colonize intracellularly host plant roots to exchange nutrients (Genre et al. 2020). The establishment of the AM symbiosis relies on the activation in root cells of the Common Symbiotic Signalling Pathway (CSSP), in which Ca^2+^ signalling plays a pivotal role (Oldroyd and Zipfel, 2017). Intriguingly, phylogenomic analyses show that clade IV MLO is present exclusively in AMF-host plant species (Bravo et al. 2016; Kusch et al. 2016; Jacott et al. 2020). However, clade IV *mlo* mutants exhibit subtle and sometimes contrasting AM phenotypes: early delayed colonization in barley, wheat and *Medicago truncatula* (Ruiz-Lozano et al. 1999; Jacott et al. 2020) versus hyper-colonization upon prolonged colonization in barley (Hilbert et al. 2020). Together, these findings suggest that clade IV MLOs act as modulators of AM development, yet their precise functions remain unresolved.

In this work, we show that *Lotus japonicus* clade IV LjMLO4 is strongly induced by AM fungi and plays a role in root development. Although *Ljmlo4* mutants displayed only mild AM phenotypes, LjMLO4 was found to be involved in the Ca^2+^-dependent regulation of primary and lateral root growth, and to facilitate Ca^2+^ transport and signalling in response to LjRALF34. These findings establish LjMLO4 as a central molecular hub that connects peptide signalling, Ca^2+^ fluxes, and root system architecture, underscoring the pivotal role of MLO proteins in integrating developmental and nutritional signals.

## Results

### *LjMLO4* has a basal expression pattern at the root tip and is upregulated in cortical cells upon colonization by AM fungi in *L. japonicus*

To identify *MLO* genes in the genome of the model legume *L. japonicus,* we exploited the BLASTp tool in LotusBase (Mun et al. 2016) querying the *L. japonicus Gifu* 1.2 proteome (Kamal et al. 2020) with known *Medicago truncatula* MLO sequences (Jacott et al. 2020). We identified 15 different *LjMLO* genes, built a phylogenetic tree via the Maximum-Likelihood algorithm with the known protein MLO sequences from *L. japonicus, A. thaliana, M. truncatula*, *Hordeum vulgare* and *Oryza sativa* (Fig. 1A) and named LjMLOs according to the belonging clade. We then analyzed the LjMLO protein sequences for previously described conserved features (Kusch et al. 2016), confirming the presence of hallmark structural elements, including seven transmembrane domains, conserved cysteine residues, and key motifs such as the C-terminal calmodulin-binding domain (Supplementary Fig. S1).

**Figure 1.**
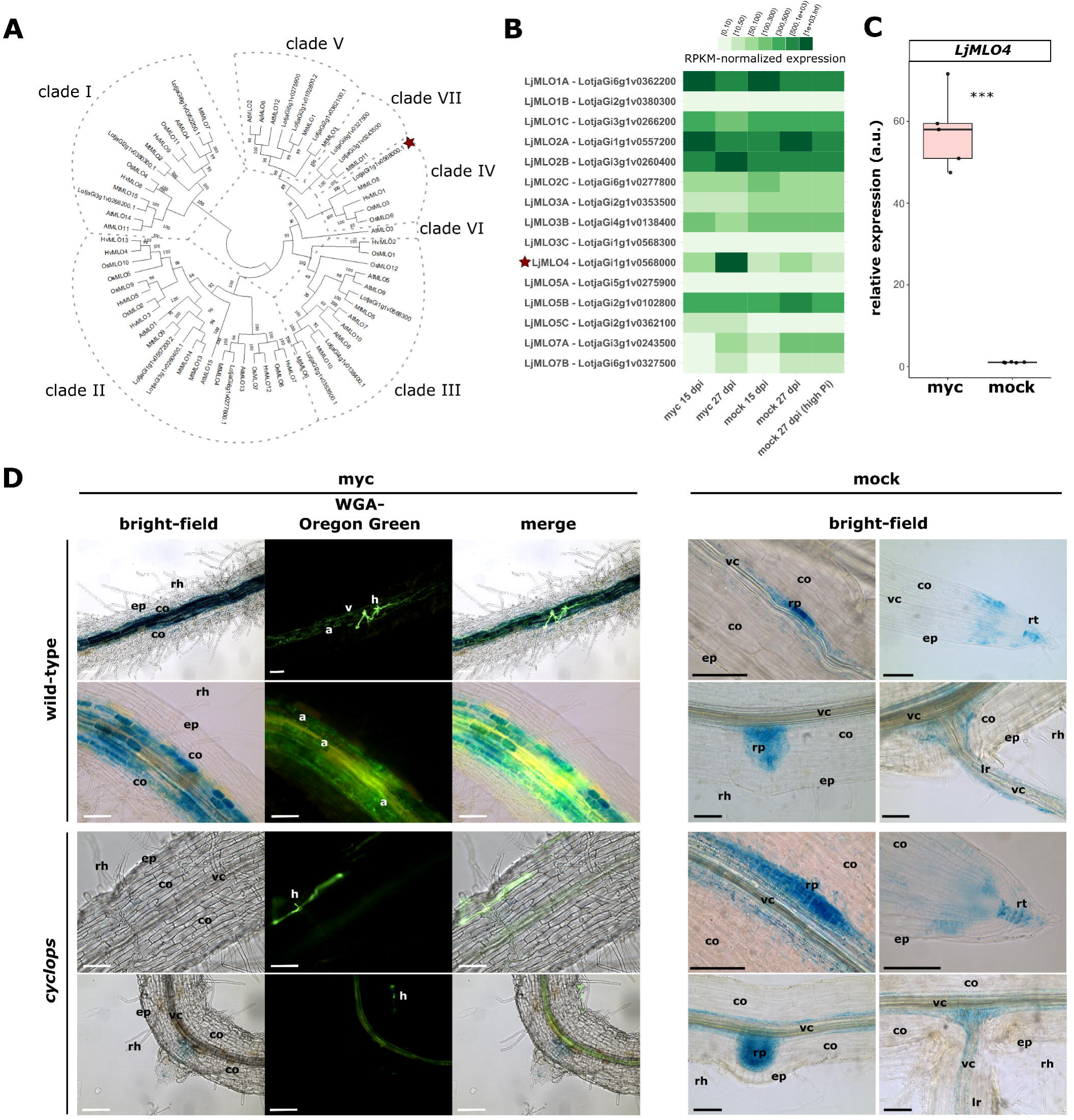
LjMLO4 is induced during arbuscular mycorrhizal symbiosis. A) Phylogenetic tree of MLO proteins from *L. japonicus*, *M. truncatula*, *A. thaliana*, *H. vulgare* and *O. sativa*. Dashed outlines indicate the major clades of the MLO family, and the red star highlights the LjMLO4 protein. Protein sequences of Mt, At, Hv, Os MLOs were retrieved from Jacott et al. (2020). LjMLO protein sequences were identified using the BLAST function in LotusBase on Gifu v1.2 genome. B) Expression pattern of *LjMLO* genes, expressed as RPKM (reads per kilobase million). Raw Expression data were retrieved from *Lotus japonicus* ExpressionAtlas in LotusBase, using the Gifu v1.2 genome dataset (Kamal et al. 2020). Rows represent the 15 *LjMLO* genes and columns are the different conditions from the dataset considered. Different shades of green represent the expression level (raw data) in the different considered conditions (darker green means higher transcript abundance). The red star highlights the *LjMLO4* gene. myc, co-cultivation with AM fungi; mock, absence of AM fungi; dpi, days post-inoculum. C) RT-qPCR analysis of the expression level of LjMLO4 in *L. japonicus* roots grown for 7 weeks in pots in the presence (myc) or absence (mock) of the AM fungus *R. irregularis*. Statistical analysis was performed using Kruskal-Wallis, followed by Dunn’s post-hoc for pairwise comparisons. ***p ≤ 0.001. D) Histochemical analysis of pMLO4::GUS activity in transformed roots of wild-type and *cyclops L. japonicus* plants after 5 weeks of co-cultivation with *R. irregularis*. Roots were stained for GUS and counterstained with WGA-Oregon Green. Representative images of mycorrhizal (myc) and non-mycorrhizal (mock) roots are shown. a, arbuscule; co, cortex; ep, epidermis; hy, hypha; lr, lateral root; rh, root hair; rp, root primordium; rt, root tip; v, vesicle; vc, vascular cylinder. Bar, 100 µm.

To investigate MLOs potential involvement in AM symbiosis, we focused on clade IV *LotjaGi1g1v0568000* (from here on *LjMLO4*) as this clade is specific to seed plants able to establish AM (Bravo et al. 2016; Jacott et al. 2021). In agreement with that, expression data retrieved from the ExpressionAtlas tool in LotusBase (Kamal et al. 2020) highlight that *LjMLO4* is extensively induced in *L. japonicus* mycorrhizal compared to control roots (Fig. 1B).

To validate the induction of the clade IV gene *LjMLO4* in mycorrhizal *L. japonicus* roots, we analyzed *LjMLO4* expression in mycorrhizal versus mock-inoculated roots and confirmed a significant 50-fold upregulation (Fig. 1C).

Then, to investigate the spatial expression pattern, its endogenous promoter (p*MLO4,* 2000 bp upstream of the start codon) was amplified from *Gifu* wild-type DNA and cloned into an expression cassette driving *UidA* expression for GUS staining. *L. japonicus* roots were transformed with *A. rhizogenes* in the wild-type and AM-defective *cyclops-3* genetic background (Yano et al. 2008) and composite plants were co-cultivated for 5 weeks in the presence (myc) or absence (mock) of the AM fungus *R. irregularis* at a low phosphate regime. *Cyclops-3* is a tilling mutant in the Gifu background mutant defective in AM and nodulation, due to the loss of the pivotal transcription factor CYCLOPS (Yano et al. 2008). After GUS staining, roots were clarified and counter-stained with WGA-Oregon Green to highlight fungal structures. In wild-type mycorrhizal roots the co-localization of the blue GUS signal with the WGA-Oregon Green signal highlights that the promoter of *LjMLO4* is strongly activated in cells colonized by AM fungal structures (Fig. 1D). Specifically, the GUS signal is mainly evident in root cortex cells and absent in epidermal cells, suggesting that LjMLO4 is likely to function in the arbuscule formation, maintenance or function. This is in line with previous microarray data coupled with laser microdissection showing that clade IV *LjMLO* is among the most highly expressed genes in root cortical cells hosting arbuscules (Guether et al. 2009) and corroborated by single cell sequencing data in tomato (Stuer et al. 2026). No fungal colonization, as well as GUS signal in the cortex, could be detected after 5 weeks of co-cultivation in the *cyclops* mutant (Fig. 1D). In addition to its induction in colonized cortical cells, p*MLO4* activity was also detected at sites of lateral root initiation and emergence, in root tips, and occasionally along the vasculature. This basal expression pattern is present in mock-inoculated roots and *cyclops* mutants, therefore independent of AM fungus presence and symbiotic signalling (Fig. 1D).

To further investigate the regulatory determinants of p*MLO4* activity, we searched for *cis-*regulatory elements (CRE) known to regulate the expression of AM-related genes (Pimprikar et al. 2018; Shi et al. 2022) and found an AW-box (WRIs binding site) and a P1BS (PHR binding site) CREs in tandem about 200 bp upstream of the start codon. Mutant promoter variants were then generated by disrupting either one CRE (ΔAW-box or ΔP1BS) or both (ΔAW-boxΔP1BS), and these constructs were tested using the GUS reporter assay. The results revealed a variable pattern of promoter activity in root cells colonized by *R. irregularis* ranging from light blue staining to the complete absence of the signal (Supplementary Fig. S2). Even in the absence of AM fungi, the promoter activity pattern of mutated versions for the AW-box and P1BS CREs did not show drastic changes (Supplementary Fig. S3). These findings suggest that both the P1BS and the AW-box CREs participate, although not critically, in regulating *LjMLO4* expression both in the presence and absence of AM fungi.

Overall, the LjMLO4 promoter exhibits a dual spatial expression pattern that depends on the colonization status of the root: basal activity mainly in actively growing root cells and strong induction in cortical cells hosting AM fungi.

### *Ljmlo4* insertional mutants are not impaired in arbuscular mycorrhiza development, but show growth penalties

Given the high induction of *LjMLO4* in roots colonized by AM fungi, we evaluated the AM phenotype in the insertional mutant lines *mlo4-1* and *mlo4-3*, isolated from the LORE1 collection (Małolepszy et al. 2016), in which the retrotransposon insertion is either in the first exon (*mlo4-1*) or in the seventh exon (*mlo4-3*) (Supplementary Fig. S4A). To determine the effect of these insertions on *LjMLO4* transcript accumulation, RNA was extracted from fully colonized roots of the wild-type, *mlo4-1* and *mlo4-3* plants. After cDNA synthesis, semi-quantitative PCR analyses targeting different regions of the *LjMLO4* transcript were performed and complemented by qRT-PCR analysis (Supplementary Fig. S4B-C). These analyses showed that *mlo4-1* roots lack detectable *LjMLO4* transcript and therefore represents a knockout line, whereas *mlo4-3* retains partial transcript accumulation, consistent with the potential production of a truncated protein, although downregulated.

Wild-type, *mlo4-1* and *mlo4-3* plants were co-cultivated with *R. irregularis* and plants were harvested after 3 and 7 weeks. Root colonization, quantified via the Trouvelot method, revealed no appreciable differences between wild-type and mutant plants (Fig. 2A and Supplementary Fig. S5) and no morphological differences in the shape of arbuscules were observed at the confocal microscope after staining with WGA-Oregon Green (Fig. 2B). Transmission electron microscopy (TEM) analyses showed a similar ultrastructural organization of colonized cells, in which the trunk and the fine branches of arbuscules are visible, in both wild-type and *mlo4* roots (Fig. 2C). In contrast, the malachite green-based quantification of soluble phosphate revealed increased phosphate accumulation in leaves of *mlo4-1* in mycorrhizal plants only, indicating that the effect is dependent on fungal colonization and likely reflects altered symbiotic phosphate nutrition (Fig. 2D). Consistent with this result, expression of *LjPT4*, a marker gene associated with arbusculated cells (Volpe et al. 2016), was significantly higher in *mlo4-1* plants (Fig. 2E). By contrast, *LjSbtM1*, another gene expressed in arbusculated cells but not directly related to phosphate uptake (Takeda et al. 2009), was downregulated in both mutant lines (Fig. 2E). Despite the absence of a strong mycorrhization phenotype in *mlo* mutants, biomass measurements showed that *mlo4* plants were generally smaller than the wild-type in terms of root, shoot and total fresh biomass, both in the presence and absence of *R. irregularis* (Fig. 2F-H). In particular, *mlo4-1* displayed a consistent reduction in biomass irrespective of mycorrhizal colonization. By contrast, *mlo4-3* plants showed a milder phenotype, which was more evident under mycorrhizal conditions. This is consistent with the more distal LORE1 insertion in *mlo4-3* and supports the possibility that this allele still produces a truncated protein retaining partial function (Supplementary Fig. S4).

**Figure 2.**
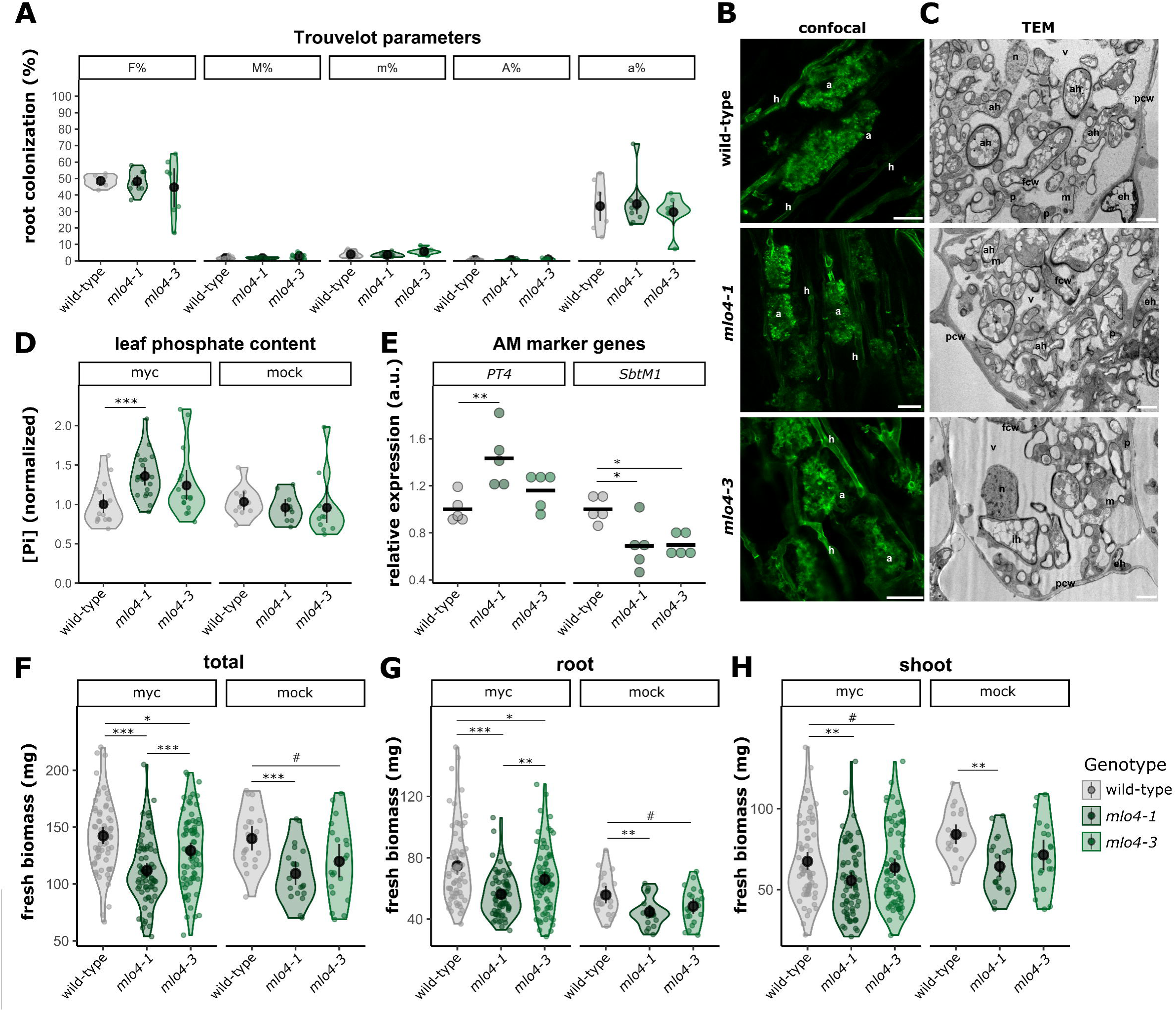
LjMLO4 is dispensable for AM colonization but affects leaf phosphate accumulation and plant growth. A) Quantification of fungal colonization in wild-type, *mlo4-1* and *mlo4-3 L. japonicus* roots after 3 weeks of co-cultivation with *R. irregularis*. Colonization parameters were calculated according to the Trouvelot method: F%, frequency of mycorrhiza; M%, absolute intensity of mycorrhiza; m%, relative intensity of mycorrhiza; A%, arbuscule abundance; a%, relative arbuscule abundance. Each dot represents one biological replicate, and data distribution is shown as violin plots, black dots indicate the median and lines indicate the confidence intervals. No significant differences were identified after statistical analysis with ANOVA. B) Confocal analysis of arbuscule morphology in colonized wild-type, *mlo4-1* and *mlo4-3 L. japonicus* roots. Roots were harvested after 7 weeks of co-cultivation with *R. irregularis* and stained with WGA-Oregon Green to visualize fungal structures. Representative images are shown. a, arbuscule; h, hypha. Bar, 20 μm. C) Transmission Electron Microscopy (TEM) observations of arbusculated cortical cells from wild-type, *mlo4-1* and *mlo4-3* roots. ah, arbuscular hypha; eh, extracellular hypha; fcw, fungal cell wall; ih, intracellular hypha; m, plant mitochondrion; n, plant nucleus; p, plastid; pcw, plant cell wall; v, plant vacuole. Bar, 2 μm. D) Quantification of soluble phosphate in leaf triplets from wild-type, *mlo4-1* and *mlo4-3 L. japonicus* plants grown in the presence (myc) or absence (mock) of *R. irregularis*. Phosphate content ([P_i_]) is expressed relative to the wild-type. Each dot represents one biological replicate, and data distribution is shown as violin plots, black dots indicate the median and lines indicate the confidence intervals. Statistical analysis was performed using Kruskal-Wallis, followed by Dunn’s post-hoc for pairwise comparisons. E) Relative expression of the AM marker genes *LjPT4* and *LjSbtM1* in roots after 7 weeks of co-cultivation with *R. irregularis*. Gene expression was quantified by RT-qPCR and normalized to the mean value of the wild-type. Each dot represents one biological replicate. Statistical analysis was performed using ANOVA, followed by Tukey’s-HSD for pairwise comparisons. F-H) Fresh biomass of the whole plants (F), roots (G) or shoots (H) of wild-type, *mlo4-1* and *mlo4-3 L. japonicus* plants grown in pots for 3 weeks with (myc) or without (mock) *R. irregularis.* Each dot represents one biological replicate, and data distribution is shown as violin plots, black dots indicate the median and lines indicate the confidence intervals. Statistical analysis was performed using Kruskal-Wallis, followed by Dunn’s post-hoc for pairwise comparisons. #p≤0.1; *p ≤ 0.05; **p ≤ 0.01; ***p ≤ 0.001.

Overall, even if the promoter of *LjMLO4* is strongly activated in mycorrhizal roots and *mlo4-1* accumulates more phosphate in the leaves than wild-type plants, no root colonization mutant phenotype could be identified in the *mlo4* plants. Instead, the reduced biomass observed also in non-colonized plants, coupled with the promoter activity detected in sites of lateral root initiation and in root tips, points to an additional role for LjMLO4 in root development beyond AM symbiosis.

### The root development phenotype of *mlo4* mutant is independent of AM colonization and modulated by external Ca^2+^

Growth penalties are a known pleiotropic effect of mutations in *MLO* loci (Acevedo-Garcia et al. 2014). To dig more into the developmental mutant phenotype of *mlo4,* we monitored root growth in axenic conditions in sterile Petri dishes. Both *mlo4-1* and *mlo4-3* mutant seedlings showed a strongly significant reduction in primary root length (Fig. 3A) and in production of lateral roots over time (Fig. 3B), further confirming that *LjMLO4* regulates root development independently of AM fungi.

**Figure 3.**
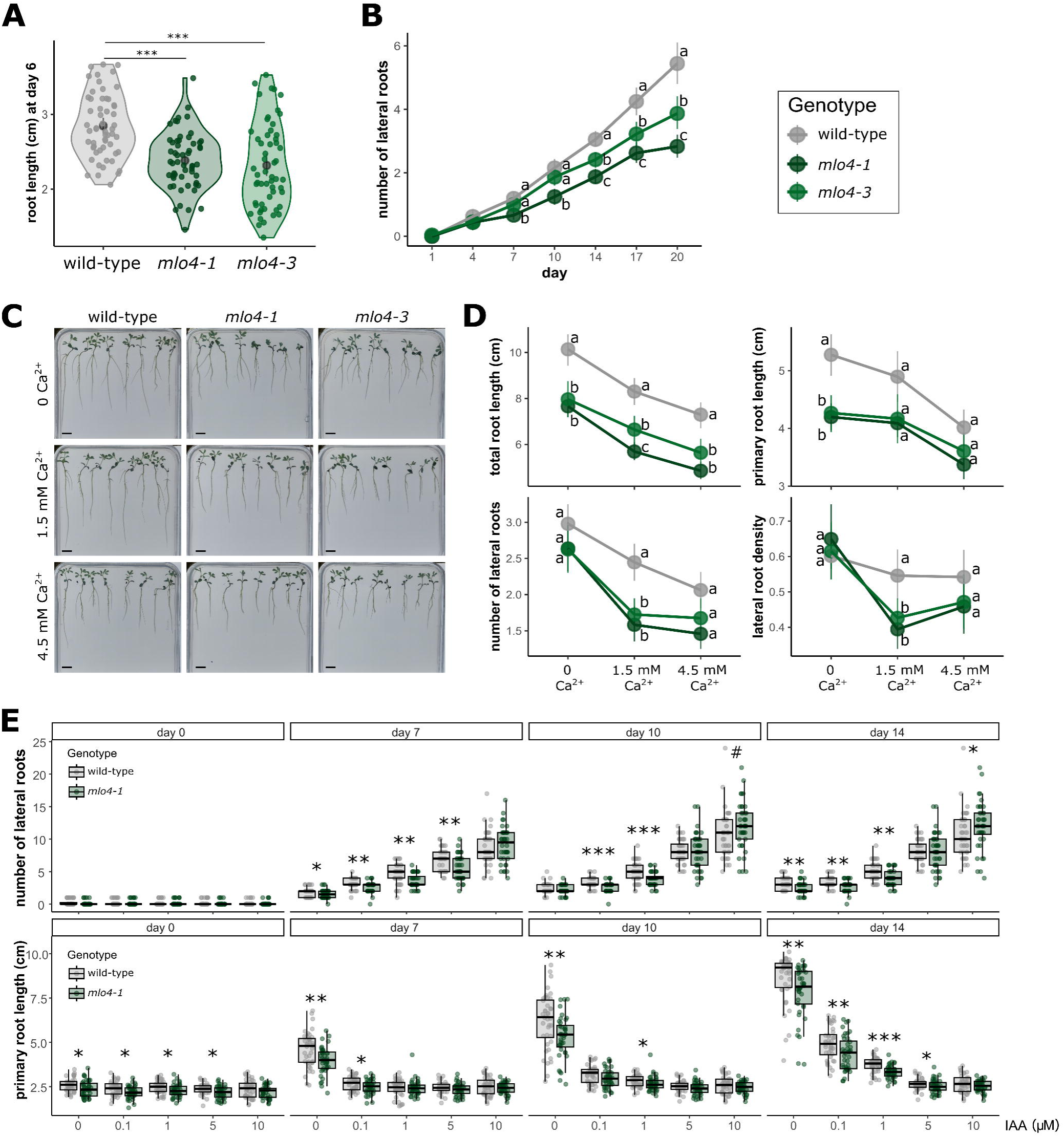
LjMLO4 influences root development independently of AM fungal colonization and in a Ca^2+^-dependent manner. A) Primary root length of wild-type, *mlo4-1* and *mlo4-3 L. japonicus* seedlings grown on square plates containing modified LA solid medium with 200 μM Pi for 6 days after seed sterilization. Each dot represents one biological replicate, and data distribution is shown as a violin plot. Statistical analysis was performed using the Kruskal-Wallis test followed by Dunn’s correction for multiple comparisons. ***p<0.001. B) Time-course analysis of lateral root formation in the absence of AM fungi. Seven-day-old wild-type, *mlo4-1* and *mlo4-3 L. japonicus* seedlings were grown on 12x12 cm square plates containing a modified LA 200 µM Pi solid medium. The number of lateral roots was counted regularly for 20 days. Data are shown as means (dots) with 95% confidence intervals (vertical lines) of the number of lateral roots for each timepoint (day). The broken line connects the dots at the different timepoints for each genotype. Statistical analysis was performed via Kruskal-Wallis test, followed by Dunn’s correction for multiple comparisons. Different letters indicate significant differences among genotypes at each time point (n≥16). C) Representative images of wild-type, *mlo4-1, mlo4-3* seedlings grown for 11 days on a modified LA solid medium containing 200 μM Pi and the indicated Ca^2+^ concentrations. Bar, 1 cm. D) Total root length, primary root length, number of lateral roots and lateral root density in wild-type, *mlo4-1* and *mlo4-3* plants grown for 11 days on modified LA solid medium containing 1% (m/v) plant agar, 200 μM Pi, and 0, 1.5 mM or 4.5 mM Ca^2+^. Statistical analysis was performed via Kruskal-Wallis test, followed by Dunn’s correction for multiple comparisons. Different letters indicate significant differences among genotypes within each Ca^2+^ treatment (n≥40). E) Number of lateral roots and primary root length in wild-type and *mlo4-1* plants grown on modified LA solid medium containing 200 μM Pi, 1% (m/v) plant agar, supplemented with the indicated IAA concentrations, and measured at 0, 7, 10, 14 days (n≥40). Statistical analysis was performed via Kruskal-Wallis test, followed by pairwise comparison with Dunn Test. #p<0.1; *p<0.05; **p<0.01; *** p<0.001.

A feature of MLO-mediated phenotypes is their modulation by exogenous Ca^2+^, as reported for clade V MLO-mediated powdery mildew infection (Bayles and Ayst 1986) and clade I MLO-mediated root thigmomorphogenesis (Chen et al. 2011; Bidzinski et al. 2014). To determine whether altered root development in *mlo4* mutants also exhibits modulation by exogenous Ca^2+^, *L. japonicus* seedlings were cultivated in an agarized medium containing varying concentrations of CaCl_2_: 1.5 mM (the standard concentration in Long-Ashton medium), 0 mM (Ca^2+^ depleted condition) and 4.5 mM (threefold enriched). Plants were imaged over time and the primary and total root length, as well as number and density of lateral roots, were monitored (Fig. 3C). All genotypes displayed the longest primary root and the highest number of lateral roots under Ca^2+^-depleted conditions (0 mM) and both parameters decreased as the external Ca^2+^ concentration increased, although with genotype-specific sensitivities (Fig. 3D). Across all [Ca^2+^] conditions, *mlo4* mutants developed a smaller root system than the wild-type as shown by the reduced total root length. However, individual root traits responded differently to changes in extracellular [Ca^2+^]. The primary root length was significantly reduced in *mlo4-1* only under 0 mM Ca^2+^ and differences in lateral root number became apparent only at 1.5 mM Ca^2+^. In both root traits, the highest [Ca^2+^] tested (4.5 mM) abolished these differences. Considering that primary root length and the number of lateral roots were differentially affected by Ca^2+^ levels, we calculated the root density (number of lateral roots/primary root length), which resulted to be affected by the exogenous [Ca^2+^] in the mutant genotypes, but not in wild-type plants. Overall, *mlo4* mutant plants displayed a distinct root developmental phenotype compared with the wild-type, and this phenotype was further modulated by exogenous Ca²⁺ concentration.

Previous reports showed that clade I MLO-mediated root thigmomorphogenesis and gravitropic response are associated with auxin signalling (Chen et al. 2011; Bidzinski et al. 2014). To test whether the root developmental phenotype of *mlo4* is influenced by auxin, wild-type and *mlo4-1* seedlings were grown on medium supplemented with increasing concentrations of indole 3-acetic acid (IAA; 0, 0.1, 1, 5 and 10 µM), and primary root length, lateral root number and root density were quantified over time (Fig. 3E and Supplementary Fig. S6). As expected, exogenous IAA strongly affected root architecture in both genotypes, in a dose-response manner. Already 7 days after treatment, IAA promoted lateral root formation, while inhibiting primary root growth, with this effect being more visible after 10 days. Importantly, the *mlo4-1* phenotype remained evident under control conditions and at low IAA concentrations: compared with the wild type, *mlo4-1* consistently formed fewer lateral roots and displayed shorter primary roots. By contrast, at higher IAA concentrations (5 and 10 µM), these genotypic differences became progressively less pronounced, indicating that strong auxin treatment tends to mask the mutant phenotype.

Together, these results indicate that *LjMLO4* plays a role in shaping root system architecture in a manner that is largely independent of AM colonization, strongly modulated by external Ca^2+^ availability and masked only upon strong exogenous auxin application.

### *mlo4* mutants show a reduced primary root growth inhibition in response to the endogenous peptide LjRALF34

MLO proteins have recently been implicated in intracellular signalling pathways activated by the plant peptide hormones Rapid Alkalinization Factors (RALFs) (Gao et al. 2022, 2023), which are known regulators of root growth and plant-microbe interactions (Blackburn et al. 2020; Abarca et al. 2021). Therefore, we tested whether LjMLO4 might also act in roots as a component of a RALF-dependent signalling pathway.

We first identified *L. japonicus* RALF peptides by BLASTp searches in LotusBase (Mun et al. 2016) using known *A. thaliana* RALF sequences as queries (Abarca et al. 2021). This search retrieved eight RALF-like peptides in the *L. japonicus* proteome. Their expression profile was obtained from the ExpressionAtlas in LotusBase (Supplementary Fig. S7A) and their protein sequences were used to build a phylogenetic tree via the Maximum-Likelihood method together with RALF sequences from *A. thaliana, M. truncatula, Solanum lycopersicum* and *O. sativa* (Fig. 4A). Based on the expression pattern, the phylogenetic tree and known biological functions of their closest *A. thaliana* homologs, we selected two candidates for functional analysis. The first, LotjaGi2g1v0402200 (hereafter LjRALF34) shows a basal expression level in roots which increases during AM colonization (Supplementary Fig. S7A) and is the closest homolog of AtRALF34 (87.5% sequence identity; Fig. 4B), a peptide involved in primary root growth and lateral root initiation (Murphy et al. 2016; Gonneau et al. 2018; Kiryushkin et al. 2023). The second, LotjaGi1g1v0770800 is highly expressed in *L. japoonicus* roots and clusters with AtRALF23 and AtRALF33 (Fig. 4A), two peptides known to negatively regulate plant immune responses (Stegmann et al. 2017) and inhibit primary root growth (Abarca et al. 2021). Given the higher sequence identity with AtRALF33 (85.7%, Fig 4C), we named this peptide LjRALF33. Both LjRALF34 and LjRALF33 are predicted to contain typical RALF domains by INTERPRO (Supplementary Fig. S7B-C).

**Figure 4.**
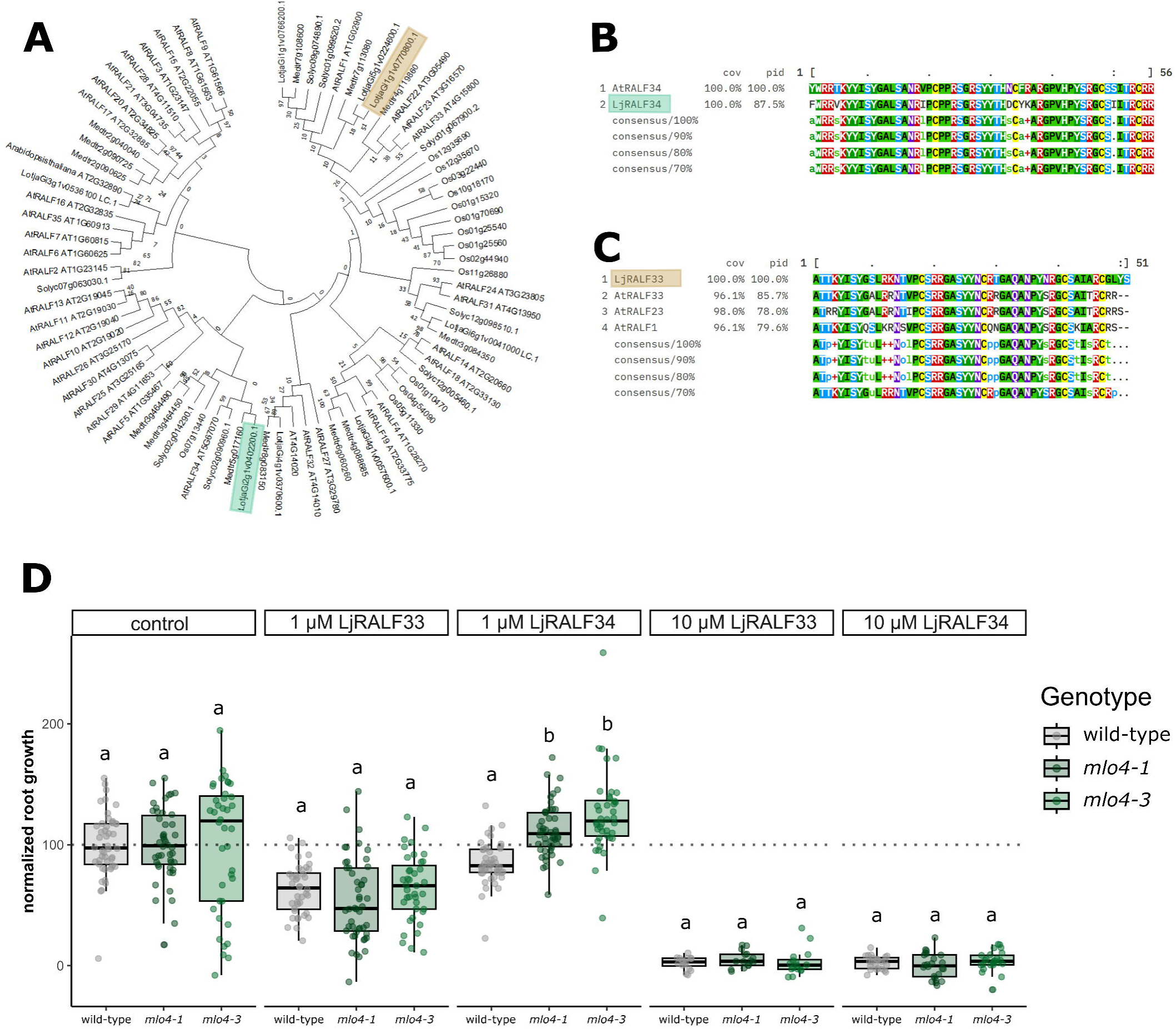
*mlo4* mutants are less sensitive to treatment with LjRALF34 peptide. A) Phylogenetic tree of *L. japonicus*, *M. truncatula*, *A. thaliana*, *S. lycopersicum*, *O. sativa* RALF peptides. *LotjaGi2g1v0402200* (LjRALF34, green) and *LotjaGi1g1v0770800* (LjRALF33, brown) are highlighted. (B) Clustal Omega alignment of the LjRALF34 and AtRALF34 peptide sequences. (C) Clustal Omega alignment of LjRALF33 with AtRALF33, AtRALF23, AtRALF22 and AtRALF1 peptide sequences. In (B) and (C) colours indicate amino acid classes based on physicochemical properties, and consensus sequences are shown below the alignments. D) Primary root growth response to LjRALF peptide treatments. Seven-day old *L. japonicus* seedlings of wild-type, *mlo4-1* and *mlo4-3 genotypes* were treated for 4 days with 1 µM or 10 µM LjRALF34 or LjRALF33, or left untreated (control). Root growth was calculated for each seedling as the difference between final and initial primary root length and is shown normalized to the corresponding control genotype. Boxplots show the median and interquartile range and each dot represents one biological replicate. Statistical analysis was performed using the Kruskal-Wallis test, followed by Dunn’s post-hoc for pairwise comparisons. Different letters indicate significantly different statistical groups within each treatment.

We then tested the effect of these peptides on primary root growth in wild-type, *mlo4-1* and *mlo4-3* seedlings. Seven-day-old plants were transferred to hydroponic culture and treated for four days with synthetic mature LjRALF34 and LjRALF33 peptides at 1 or 10 µM, or water (control condition). Primary root length was measured before treatment (day 0) and again after four days of treatment. Because the genotypes differed in root length at day 0, root growth after treatment was expressed as percentage growth inhibition relative to the corresponding untreated control of each genotype (Fig. 4D and Supplementary Fig. S8). Although both *LjRALF33* and *LjRALF34* at 10 µM completely inhibited root growth in all three genotypes, their effects differed at lower concentrations. Treatment with 1 µM LjRALF33 caused a similar reduction in root growth in all genotypes, corresponding to approximately 80% inhibition. By contrast, whereas 1 µM *LjRALF34* mildly inhibited root growth in wild-type plants, both *mlo4* mutant lines showed higher normalized root growth, indicating a reduced sensitivity to this peptide.

Overall, these results indicate that *mlo4* mutants are specifically less sensitive to *LjRALF34*-mediated inhibition of primary root growth, supporting a role for *LjMLO4* in the signalling pathway activated by this peptide.

### The heterologous expression of LjMLO4 in *E. coli* facilitates Ca^2+^ influx

Our results support the existence of a RALF-MLO signalling module regulating various aspects of plant development, as previously reported in *A. thaliana* (Gao et al. 2022, 2023). MLOs have been shown to be cation-permeable channels and to be involved in the generation of the Ca^2+^ signals triggered by RALFs perception (Gao et al. 2022, 2023). To examine if this is also the case for LjMLO4, we evaluated its ability to mobilize Ca^2+^ across membranes by heterologous expression in *E. coli* cells. To facilitate translation, the coding sequence of *LjMLO4,* tagged with polyhistidine (6xHis) at the C-terminal region, was codon-optimized for the expression in the bacterial system and cloned into a pRham vector for rhamnose-dependent induction. Along with the full-length protein LjMLO4, a truncated version (AA 1-265, LjMLO4Δ) was cloned to exclude the FWF predicted ion transport motif (Kusch et al. 2016) and the C-terminal calmodulin-binding motif, mimicking the putative protein product of the *mlo4-3* mutant insertion line. Different versions of the pRham expression vector (empty, MLO4, MLO4Δ) were then co-expressed with pACYCDuet-1(HA-Aeq), encoding the Ca^2+^-sensitive photoprotein aequorin (Teardo et al. 2019). Upon induction of protein expression, the transformed *E. coli* cells were exposed to external Ca^2+^ pulses (1 mM CaCl_2_), and the resulting changes in intracellular [Ca^2+^] were monitored using aequorin-based Ca^2+^ measurement assays. The expression of both LjMLO4 and LjMLO4Δ was confirmed by Western blotting with an anti-His antibody (Supplementary Fig. S9A) and the correct functionality of the Ca^2+^ reporter was validated by measurements of the total luminescence emitted upon Ca^2+^ discharge (Supplementary Fig. S9B). All bacterial strains showed high levels of luminescence (>10^6^ photons), allowing a reliable quantification of intracellular Ca^2+^ levels (Ottolini et al. 2014). As shown in Fig. 5A-D, the exogenous Ca^2+^ pulse triggered a Ca^2+^ influx characterized by different dynamics over time when the full length LjMLO4 is expressed compared to the empty vector and LjMLO4Δ. Despite a similar and immediate Ca^2+^ peak (Fig. 5C-D), LjMLO4-expressing cells mobilize more Ca^2+^ over time (Fig. 5D). When the control solution was applied to the bacterial cells, only the physiological Ca^2+^ response to touch (Moscatiello et al. 2010) was detected, and no differences were observed between LjMLO4, LjMLO4Δ, and the empty vector (Supplementary Fig. S9C-F).

**Figure 5.**
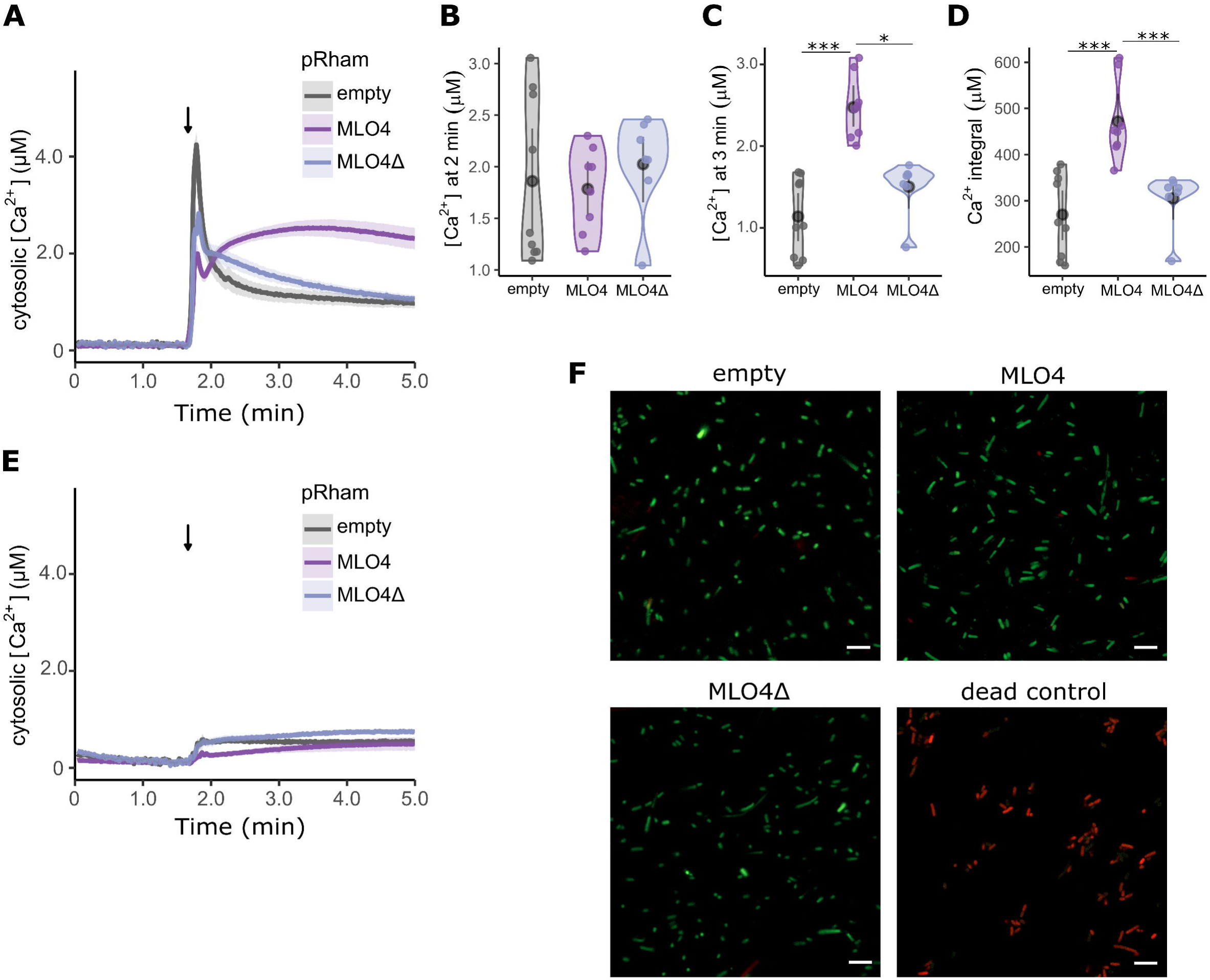
Heterologous expression of LjMLO4 enhances Ca^2+^ influx in aequorin-expressing *E. coli* cells. *E. coli* cells were co-transformed with pET-AEQ for aequorin expression and with one of the pRham constructs: empty control vector (empty, grey), codon-optimized LjMLO4 CDS (MLO4, purple), or a truncated LjMLO4 version (MLO4Δ, blue) mimicking the putative truncated protein in *mlo4-3 L. japonicus* mutant line. A-D) Cytosolic Ca^2+^ changes in response to 1 mM CaCl_2_. A) Ca^2+^ concentration over time presented as means ± SE (shading); the arrow indicates the time of CaCl_2_ addition. B,C) Ca^2+^ concentration measured 2 min (B) and 3 min (C) after the external Ca^2+^ pulse. D) Integrated Ca^2+^ dynamics over time (5 min). Each dot represents an independent biological replicate: distributions are shown as violin plots, black dots indicate the median and lines indicate the confidence intervals. Statistical analysis was performed using the Kruskal-Wallis test followed by Dunn’s post-hoc correction. *p<0.05; *** p<0.001. E) Ca^2+^ traces in response to 1 mM CaCl_2_ after 10 min pre-treatment with the Ca^2+^ channel inhibitor LaCl_3_; data are presented as means ± SE (shading) F) Viability of transformed *E. coli* cells. Green (SYTO 9) indicates live cells, red (propidium iodide) indicates dead cells. Cells incubated at 100°C for 10 min were used as dead control. Bar, 5 μm.

Pre-treatment with the ion channel blocker LaCl_3_ effectively abolished the transient increases in cytosolic Ca^2+^ both in the presence and absence of LjMLO4 (Fig. 5E and Supplementary Fig. S9G), supporting the specificity of the observed Ca^2+^ fluxes.

The viability of the aequorin-expressing *E. coli* cells was not affected by LjMLO4 or LjMLO4Δ presence (Fig. 5F). The expected localization of both versions of the protein at the periphery of bacterial cells was verified by immunofluorescence analyses (Supplementary Fig. S9H) and the preservation of an unaltered ultrastructure was confirmed by TEM observations (Supplementary Fig. S9I).

Altogether, these findings strongly point to LjMLO4 functioning as a Ca^2+^-permeable channel capable of mediating Ca^2+^ fluxes across cellular membranes, similar to other MLOs, such as HvMLO in barley and clade V AtMLO2/AtMLO12 (Gao et al. 2022). On the other hand, the truncated version of the protein (LjMLO4Δ) was found to be altered in this function.

### LjRALF34 peptide triggers root zone-specific Ca^2+^ transients and *Ljmlo4* insertional mutants show a reduced Ca^2+^response at the root apex

After showing that *mlo4* mutant roots are less sensitive to LjRALF34 treatment and that LjMLO4 has the ability to mobilize Ca^2+^ across membranes in *E. coli* experiments, we tested whether LjMLO4 mediates responses to LjRALF34 by activating a Ca^2+^ signalling cascade. To this aim, *L. japonicus* roots were transformed via *A. rhizogenes-*mediated hairy root transformation to express a cytosol-only localized YFP-aequorin (Binci et al. 2024). Transformed root segments were treated with the RALF synthetic peptides and the cytosolic [Ca^2+^] was monitored over a 30 min time interval. Intriguingly, we found that LjRALF34 triggered distinct Ca^2+^ dynamics depending on the distance from the root apex (Supplementary Fig. S10A-F). Apex-containing root segments responded with a steep and intense Ca^2+^ elevation a few seconds after the application of the stimulus, which was much less pronounced in root segments distal from the root apex. After a fast decrease, [Ca^2+^] remains higher than the basal level over time in both root zones. Different doses of the peptide (1 and 10 µM) triggered Ca^2+^ transients characterized by a similar pattern, but with different magnitudes of the Ca^2+^ peak. Similar Ca^2+^ responses were triggered by LjRALF33 (Supplementary Fig. S11A-C).

We next examined if LjMLO4 is involved in the generation of the RALF-induced Ca^2+^ signals. To this aim, Ca^2+^ dynamics in *mlo4-1* and *mlo4-3* root segments-expressing cytosolic YFP-aequorin were analyzed in comparison with the wild-type. Both *mlo4* insertional mutant lines responded with Ca^2+^ transients with dynamics similar to the wild-type in root segments derived from different root zones. Whereas no significant differences among genotypes were visible after treatment with 10 µM LjRALF34 (Fig. 6A-C and Supplementary Fig. S10G-I), both *mlo4* mutants responded with reduced maximal [Ca^2+^] elevation and overall Ca^2+^ mobilization to 1 µM LjRALF34 (Fig. 6D-F) compared to the wild-type. This result is in line with the reduced sensitivity of *mlo4* mutant roots to LjRALF34-triggered growth inhibition. On the other hand, no clear differences in the Ca^2+^ traces could be found in response to LjRALF33 (Supplementary Fig. S11D-I). Considering the strong induction of LjMLO4 during AM symbiosis, and the broader role of MLOs in plant-microbe interactions, we also tested if *Ljmlo4* roots-expressing cytosolic YFP-aequorin were less sensitive to a range of abiotic and biotic stimuli. In response to salt stress and oxidative stress, as well as short-chain (CO4) and long-chain (CO7) chitin oligosaccharides, no significant differences in Ca^2+^ dynamics emerged among the genetic backgrounds, with the only exception of the CO7-treated *mlo4-3* (Supplementary Fig. S12).

**Figure 6.**
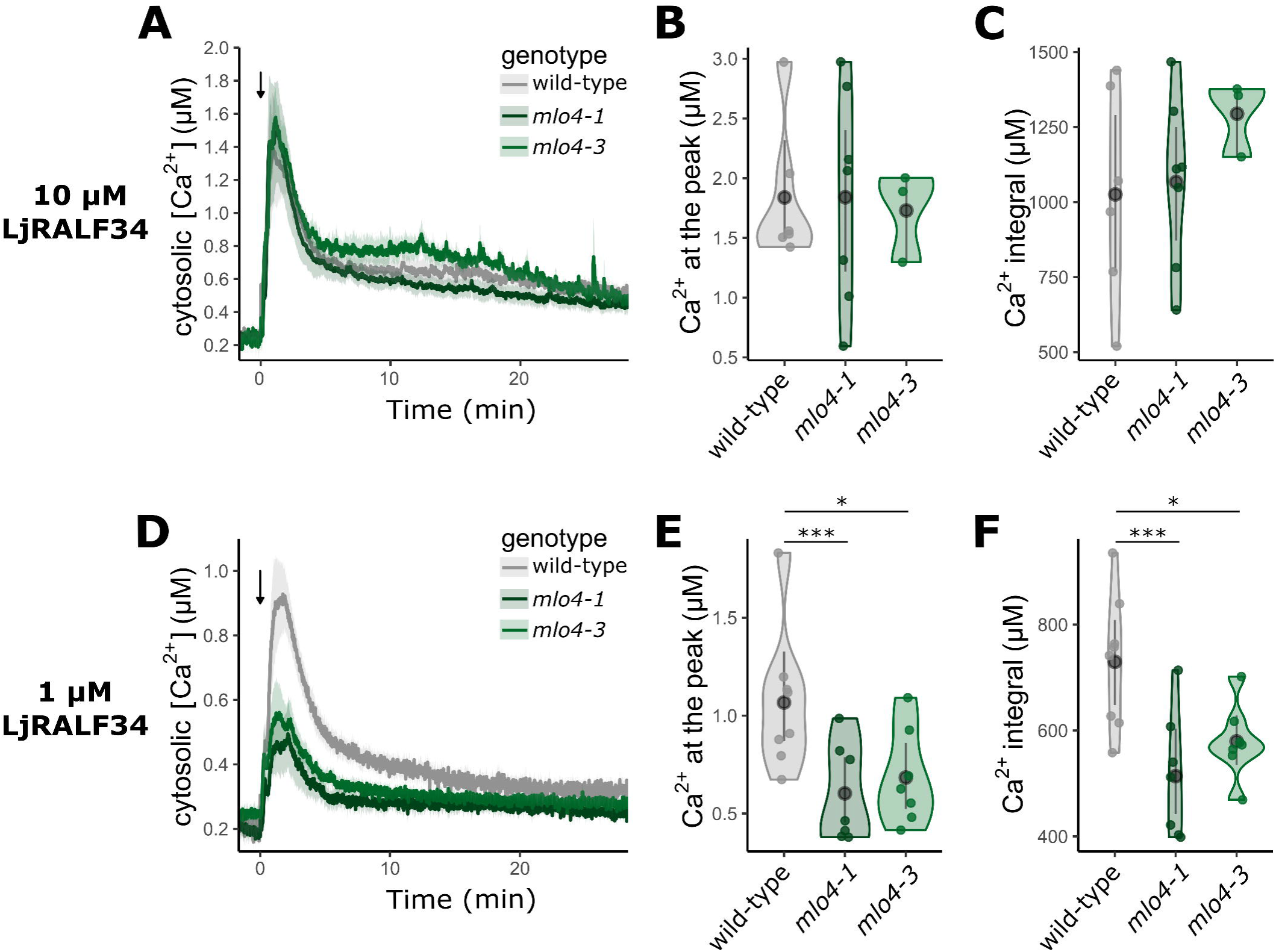
*mlo4* mutants show a reduced Ca^2+^ response to 1 µM LjRALF34 at the root apex. Cytosolic Ca^2+^ changes triggered by 10 µM and 1 µM LjRALF34 were monitored in root apex-containing segments of *L. japonicus* wild-type, *mlo4-1, mlo4-3* plants expressing cytosolic YFP-aequorin. A, D) Data are presented as means ± SE (shading) of n ≥ 3 obtained from at least 3 different composite plants (independent transformations). The stimulus was injected at time 0 (min), indicated by the arrow. B, E) Dots represent the maximum [Ca^2+^] for each run. C, F) Integrated Ca^2+^ dynamics over time (30 min). Distributions are shown as violin plots, black dots indicate the median and lines indicate the confidence intervals. Statistical analysis was conducted via Kruskal-Wallis test followed by Dunn’s post-hoc correction. *p<0.05; *** p<0.001.

To sum up, we found that LjMLO4 is a mediator of the Ca^2+^-mediated LjRALF34-triggered intracellular signalling cascade in roots, further supporting that the LjRALF34-LjMLO4 module is a regulator of root development in *L. japonicus*.

## Discussion

### Clade IV MLOs are involved in AM symbiosis, but their precise role remains elusive

In this work, we analyzed the role of clade IV LjMLO4 in *L. japonicus*, whose homologs in barley and *M. truncatula* are known to regulate colonization by filamentous fungal pathogens and mutualists (Büschges et al. 1997; Ruiz-Lozano et al. 1999; Hilbert et al. 2020; Jacott et al. 2020). *LjMLO4* was strongly upregulated in cortical cells colonized by the AM fungus *R. irregularis*, yet *mlo4* mutants displayed no obvious defects in root colonization. At the same time, mycorrhizal *mlo4* plants accumulated more phosphate in leaves and showed increased expression of the AM-associated phosphate transporter LjPT4, suggesting that nutrient exchange or arbuscule lifespan may be enhanced in the absence of *LjMLO4*. These findings add further complexity to previous reports on clade IV *MLO* silencing in different species, which described either delayed early colonization or increased colonization at later stages (Hilbert et al. 2020; Jacott et al. 2020). Such variability in colonization phenotypes could arise from technical differences, including host plant and fungal species, quantification methods, or growth conditions, such as nitrogen regimes and light intensity, which are known to affect *mlo*-related pleiotropic phenotypes (Freh et al. 2024). In addition, the expression of multiple *MLO* genes in mycorrhizal roots (Jacott et al. 2020) suggests potential functional redundancy, which may mask the specific contribution of clade IV MLO to AM symbiosis. As reported for powdery mildew resistance in Arabidopsis (Consonni et al. 2006), higher-order *mlo* mutants may therefore be required to uncover more severe symbiotic phenotypes. Another crucial factor is the complexity of AM development, characterized by different stages in which MLOs may exert their biochemical function. Increasing evidence suggests that MLO proteins function as Ca^2+^-permeable channels (Gao et al. 2022, 2023; Li and Xiao 2025; Hübbers et al. 2026). In agreement with this, heterologous expression of LjMLO4 in *E. coli* was found to significantly enhance Ca^2+^ mobilization across bacterial membranes. The specificity of LjMLO4-dependent Ca^2+^ transport was validated via pharmacological manipulation of Ca^2+^ homeostasis, viability assay and TEM and confocal imaging, excluding alterations of the bacterial membrane.

MLO protein features hint at a possible involvement in plant-fungus communication before and during infection, although our data argue against a major role for LjMLO4 in the Ca^2+^-mediated perception of chitin-based fungal signals (Giovannetti et al. 2024a,b). Alternatively, because MLO proteins have been linked to vesicle trafficking and cell wall composition (Hübbers et al. 2024), LjMLO4 may contribute to membrane remodelling at the plant-fungus interface or to cell wall dynamics during fungal penetration. The increased phosphate accumulation in *mlo4* plants, together with *LjPT4* upregulation, further suggests the intriguing possibility that MLO functions directly at the periarbuscular membrane, either directly as a Ca^2+^ channel or more indirectly as a regulator of membrane dynamics. In line with this, Alphafold 3-modelling predicted that homotrimerization of MLOs creates a transmembrane pore that can open with increased membrane tension (Hübbers et al. 2026), making it possible that MLOs act as mechanosensor during root colonization by the fungus. Finally, the contribution of LjMLO4 to AM symbiosis may be at least partly indirect, arising from its broader role in root development.

### Modulation of the root system architecture is a new function for the MLO gene family

Loss of MLO function is classically associated with resistance to powdery mildew, often accompanied by pleiotropic growth defects (Acevedo-Garcia et al. 2014). In line with this, we found that *mlo4* mutant plants show reduced root biomass, shorter primary roots and fewer lateral roots. This root developmental mutant phenotype, along with the specific activity of p*MLO4* at root tips and lateral root initiation sites, is independent from AM fungal colonization, directly implicating LjMLO4 in root system architecture. Notably, the mutant phenotype was modulated by exogenous Ca^2+^, with distinct effects on primary root elongation and lateral root formation, suggesting that LjMLO4 integrates root development with external Ca^2+^ availability. Ca^2+^ homeostasis and exogenous Ca^2+^ levels have been shown to critically balance the trade-off between growth and immunity in *A. thaliana* (Wang et al. 2024). Overall, our data provide first evidence on the function of clade IV MLO as modulator of the root system architecture in *L. japonicus,* impacting both on root elongation and lateral root formation. This function expands the repertoire of MLO activities in root biology, complementing previous reports of roles in root thigmotropism and gravitropism for clade I (Chen et al. 2009; Bidzinski et al. 2014; Zhu et al. 2021; Zhang et al. 2022) and root hair elongation for clade II MLOs (Ogawa et al. 2025). Although our results indicate that LjMLO4 shapes the development of the root independently from fungal colonization, it is likely that there is an interplay of the two functions. Indeed, chitin signalling and mycorrhizal colonization are known to affect root architecture (Chiu et al. 2022) and our findings on the biomass of colonized plants showed that growth penalties in *mlo4* are exacerbated by the fungal presence, highlighting LjMLO4 as a molecular link between root growth and symbiotic performance.

### RALF-responsive Ca^2+^ signalling may represent a conserved mode of MLO action

In this work we show that LjMLO4 can mobilize Ca^2+^ across membranes when expressed in *E. coli*, and that *L. japonicus mlo4* mutants are less sensitive than the wild-type to LjRALF34, both in terms of Ca^2+^ signalling and primary root growth inhibition. By contrast, no differences between *mlo4* and wild-type were detected in response to LjRALF33, indicating that this phenotype is specific to LjRALF34. Although both peptides are expressed in roots, their functions may be cell-type specific, consistent with the highly restricted expression pattern of LjMLO4 in non-colonized roots. This is further supported by the different Ca^2+^ transients triggered by the peptides in distinct root zones. Moreover, the two peptides in *A. thaliana* are recognized by different CrRLK1L receptors, with AtRALF23/33 acting through FERONIA (Stegmann et al. 2017) and AtRALF34 through THESEUS (Gonneau et al. 2018). Various reports point to MLO functioning downstream of CrRLK1L signaling. In synergid cells, pollen tube-derived RALF4/RALF19 bind to FERONIA, which then interacts with clade III MLO7/NORTIA activating its Ca^2+^ channel function (Gao et al. 2022). In root hairs, clade II MLO15 can rescue *feronia* defects in Ca^2+^ oscillations and tip growth (Ogawa et al. 2025), a process known to involve RALF22 (Schoenaers et al. 2024). In pollen tubes, where FERONIA is not expressed, clade II-III AtMLO1/5/9/15 function downstream of RALF and ANXUR1/2, other members of the CrRLK1L receptor family. Overall, these findings support the existence of a broad RALF/CrRLK1L/MLO signalling module operating in multiple developmental contexts (Ogawa and Kessler, 2023). Our findings extend this framework to the control of root system architecture in *L. japonicus* and support a model in which LjMLO4 enhances LjRALF34-triggered signalling in root cells. At the same time, since little is known about the CrRLK1L family in *L. japonicus*, the upstream components that perceive LjRALF34 and activate LjMLO4 remain to be identified. Finally, considering that MLOs can shape cell wall composition (Hübbers et al. 2024) and that RALFs act as sensor of the cell wall status (Moussu et al. 2023, Rößling et al. 2024), it cannot be ruled out that the reduced sensitivity of *mlo4* mutants to LjRALF34 may also reflect altered cell wall properties.

Given the apparent multifaceted role of LjMLO4, it is tempting to speculate that RALF/MLO signalling is also involved in AM symbiosis. RALFs are mainly recognized as immunity regulators and RALF-like peptides are secreted by several pathogens, such as *Fusarium* and nematodes, to suppress plant immunity (Ascurra and Müller, 2025). In addition, an endogenous RALF has been shown to negatively regulate rhizobial nodulation in *M. truncatula* (Combier et al. 2008), whereas the involvement of RALFs in AM symbiosis has not yet been investigated. Future studies are needed to explore whether endogenous RALFs participate in AM development, and if AM fungi themselves produce RALF-like peptides to modulate host signalling.

## Materials and Methods

### Protein sequence alignment, structure prediction, phylogenetic analyses and peptide synthesis

The MLO and RALF protein sequences of *L. japonicus* (Supplementary Table S1) were retrieved from LotusBase proteome *Gifu* v1.2 (Mun et al. 2016; Kamal et al. 2020). The ClustalW algorithm was used to perform the multisequence alignment with MLO and RALF sequences (Jacott et al. 2020; Campbell et al. 2017). The phylogenetic tree was built using the Maximum-Likelihood algorithm with the MegaX software, with default settings and bootstrap 1000. MLO multisequence alignment was visualized using ESPript (Robert and Gouet 2014), and LjMLO4 structure prediction was performed through the Swiss-Model homology-modelling server (Waterhouse et al. 2018).The predicted mature peptides of LjRALF33 and of LjRALF34 sequences (Supplementary Table S2) were used to synthesize the two peptides (Proteogenix, Schiltigheim, France), which were dissolved in H_2_O water, frozen in liquid N_2_ and stored at -80° C in small aliquots until use, as previously described for synthetic AtRALF peptides (Somoza et al. 2024).

### Molecular cloning and bacterial transformation

The amplification from *L. japonicus* (*Gifu* ecotype) genomic of the 2 kb promoter of *LjMLO4* and the genomic locus of *LjMLO4* was performed with Q5 DNA polymerase (New England Biolabs (NEB), Ipswich, MA, USA) according to manufacturer’s instructions. Primers (Supplementary Table S3) for promoter amplification and mutagenesis were designed in benchling.com to be compatible with the GreenGate cloning system (Lampropoulos et al. 2013). Entry and expression vectors were assembled via cut-ligation with BsaI (NEB) and T4-DNA ligase (NEB), following the GreenGate protocol.

For heterologous expression in *Escherichia coli* cells, the coding sequence of *LjMLO4* was codon optimized (Supplementary Table S2) and synthesized by Twist Bioscience’s (San Francisco, CA, USA). The optimized gene was delivered as linear gene fragments and cloned into pRham (Lucigen Corporation, Middleton, WI, USA) via homologous recombination-based cloning, according to Jacobus and Gross (2015). The same system was exploited to obtain the LjMLO4Δ-encoding plasmid.

The selection and amplification of vectors were performed in DH5α *E. coli* cells upon transformation with heat shock of chemo-competent cells. Plasmid sequences were checked via Sanger sequencing at BMR Genomics (Padova, Italy) or Eurofins Genomics (Ebersberg, Germany) and by Primordium long-read DNA sequencing at Primordium Labs (Arcadia, CA, USA) or Whole-Plasmid Sequencing (Eurofins Genomics). The expression vectors were then transformed into *Agrobacterium rhizogenes* 1193 via the freeze & thaw method (Wise et al. 2006). All plasmids used in this work are listed in Supplementary Table S4.

### Plant lines and genotyping

Wild-type *Gifu*-ecotype plants of *L. japonicus*, *cyclops-3* TILLING mutant (Yano et al. 2008) and three LORE1 mutant lines (Malolepszy et al. 2016) for the *LotjaGi1g1v0568000 LiMLO4* locus (30015501 - *mlo4-1* and 30107425 - *mlo4-3*) were used in this work. Segregant populations of the different LORE1 lines were ordered from https://lotus.au.dk/ (Mun *et al*. 2016) and genotyped to identify homozygous mutants, following the suggested primers and protocols. The same primers were used for analysis of the transcribed mRNA from the mutant *mlo4* loci.

### RNA extraction, cDNA synthesis and quantitative PCR

Total RNA was extracted from *L. japonicus* roots, after 7 weeks cultivation in pots in presence or absence of *R. irregularis*. Roots were homogenized using mortar and pestle and 250-300 mg root powder were used for RNA extraction with the RNA Plant and Fungi kit (MACHEREY-NAGEL GmbH & Co. KG, Dueren, Germany), according to the manufacturer’s instructions. cDNA synthesis was performed as previously described (Binci et al. 2024). Genotyping primers (described above) were used in a semi-quantitative RT-PCR (30 cycles) to validate genotyping at the genomic level. Real Time quantitative PCR (RT-qPCR) was performed on cDNA to evaluate the expression of *LjMLO4* and AM marker genes using the HOT FIREPOL EvaGreen qPCR mix plus (Solis BioDyne, Tartu, Estonia) on a 7500 real-time PCR system (Thermo Fisher, Waltham, MA, USA). The *LjUbiquitin10* and *LjATPsynthase* genes were used as an internal reference for analysis of the target gene expression.

### Sterilization and germination of *L. japonicus* seeds

*L. japonicus* seeds scarification and sterilization was performed as previously described (Binci et al. 2024). According to the experiment to be performed, seeds were left germinating in different setups. For mycorrhization and hairy root transformation, seeds were plated in ½ B5 (Duchefa, Haarlem, The Netherlands) medium with 1% (w/v) Plant Agar (Duchefa) in 12x12 cm squared Petri dishes and vertically incubated in the growth chambers. For phenotyping of root traits, the seeds were distributed in round Petri dishes supplied with a fine layer of H_2_O and 1% Plant Agar and incubated upside down to favour homogeneity among germinating roots. In any case, seed germination was performed in dark conditions for three days (temperature 23°C, humidity 50%). After three days, seedlings aimed for mycorrhization and hairy root transformation were exposed to light (16h/8h light/dark) and allowed to grow vertically in the same plate until use. Seedlings aimed for phenotyping of root traits were moved to a new squared Petri dish 12x12 cm supplied with a modified Long Ashton (LA) solution with 200 µM KH_2_PO_4_ (Pi) + 1% Plant Agar and grown vertically until the appropriate experiment was performed.

### Mycorrhization

15 days old *L. japonicus* seedlings were moved to plastic pots (9 cm diameter, 4 plants per pot) filled with river sand, previously washed twice with ddH_2_O and autoclaved. Propagules of *R. irregularis* (Agronutrition, Carbonee, France) were washed with ½ tap water and ½ ddH_2_O and evenly distributed within each pot (3000 spores/pot). Mock-treated plants lacked the AMF propagules. Plants were grown in growth chambers (22°C, 60% humidity, 16h/8h light-dark cycle) and fertirrigated twice per week with 20 ml liquid modified LA 20 μM Pi until use.

### Staining of AM fungi in colonized roots

For quantification of mycorrhizal structure, roots were clarified and ink-stained in an acidic solution as previously described (Binci et al. 2025). Quantification was performed according to the Trouvelot method (Trouvelot et al. 1986) using a Leica DM1000 optical microscope (Leica, Wetzlar, Germany). To analyse the morphology of arbuscules and to counterstain GUS-stained roots, roots were clarified in 10% (w/v) KOH for 3 days at room temperature, washed in PBS and incubated for 4 h in 0.1 M HCl. After two washes in PBS, roots were incubated overnight (4°C) with 1 µg/ml WGA-Oregon Green 488 (Thermo Fisher) in PBS. Images were acquired with a Zeiss LSM900 Airyscan2 (Zeiss, Oberkochen, Germany) confocal microscope using the Argon488 laser (emission 495-530 nm).

### Transmission electron microscopy (TEM) analyses

Roots from wild-type, *mlo4-1* and *mlo4-3* plants were harvested after co-cultivation with *R. irregularis* for 7 weeks. Hyphopodia were identified with the help of a stereomicroscope and root segments were excised around them. Fresh root samples were incubated in 1 mL of fixing solution (3% glutaraldehyde in 0.1 M cacodylate buffer, pH 7.4) overnight at 4°C. Washing, fixation with osmium, embedding and slicing were performed as previously described (Montanari et al. 2025). Observations were carried out with a Tecnai G2 transmission electron microscope (Field Electron and Ion Company, Hillsboro, OR, U.S.A.) operating at 100 kV and equipped with an Osis Veleta camera (Olympus, Tokyo, Japan).

### Measurement of soluble phosphate in *L. japonicus* leaf samples

The second leaf triplets from the top of each plant shoot were collected while harvesting the plant for the mycorrhization assay. Soluble phosphates were extracted as previously described (Giovannetti et al. 2019) and then quantified using the Malachite Green Phosphate Assay Kit (Merck, Darmstadt, Germany), following the manufacturer’s instructions. Absorbance data were converted into concentrations of phosphate using the calibration curve and standardized to the fresh weight of the leaf sample.

### *A. rhizogenes*-mediated hairy root transformation

Hairy root transformation was performed as previously described (Binci et al. 2024). Briefly, *L. japonicus* composite plants were generated by infecting detached shoots from 7 days old seedlings with *A. rhizogenes* AR1193 strains carrying the plasmid of interest. After 6 days of co-cultivation, shoots were grown for 4-5 weeks in solid ½ B5 with 300 µg/ml cefotaxime (Duchefa). Transformed roots were identified with the fluorescence stereomicroscope MZ16f (Leica) according to the used transformation marker.

### GUS staining

Composite plants were grown in pots containing *R. irregularis* propagules (myc) or not (mock) and watered with modified LA (20 µM Pi). Harvested roots were selected at the fluorescence stereomicroscope MZ16f (Leica) to identify transformed roots. The selected roots were incubated overnight at 37°C in the following reaction mix: 5 mM K_3_Fe(CN)_6_, 5 mM K_4_Fe(CN)_6_6H_2_O, 0.1 M potassium phosphate buffer, 1 mM Na_2_EDTA, 1% Triton X-100, 1.25 mg/ml X-Gluc (5-bromo-4-chloro-3-indolyl-beta-D-glucoronide). After the incubation time had elapsed, samples were washed in 70% (v/v) ethanol and stored in the same solvent at 4°C until use. The counterstaining with WGA-Oregon Green 488 was performed as described above. Images were then acquired with the DM6b optical microscope (Leica) in brightfield and using the I3 filter cube (Ex BP 450-490; dichroic 510; Em LP 515), at 10x, 20x and 40x magnification.

### Phenotyping of root traits

To monitor primary root growth and the emergence of lateral roots, synchronized seedlings were placed into 12x12 cm square Petri dishes containing 60 ml of modified LA 200 μM Pi with 1% Plant Agar. In each plate, 8 seedlings were evenly distributed in a straight line 3 cm away from the top side of the plate. For exogenous Ca^2+^ treatments, seedlings were placed in modified LA 200 μM Pi with 3 mM KNO3 to replace 1.5 mM Ca(NO3)_2_ added with the following concentration of CaCl_2_ and KCl to modulate Ca^2+^ levels and compensate for chloride variations: 0 CaCl_2_ and 9 mM KCl (0 Ca^2+^); 1.5 mM CaCl_2_ and 6 mM KCl (1.5 mM Ca^2+^); 4.5 mM CaCl_2_ (4.5 mM Ca^2+^). For auxin treatment, different concentrations of indole-3-acetic acid (IAA, 0.1 µM, 1 µM, 5 µM, 10 µM) or 10^-4^ M dimethyl sulfoxide (solvent control) were added directly to the growth medium prior to solidification. In both cases, seedlings were grown vertically for 2-3 weeks (23°C, humidity 50%, 16h/8h light/dark) and plates were routinely imaged with a 2D scanner.

For LjRALF treatment, seedlings were grown for 4 days after germination in square Petri dishes with solid 60 ml of LA 200 μM Pi. Seedlings were then moved to 8-strips (1.2 ml capacity per tube) putting one plant per tube containing 1 ml of liquid LA 200 μM Pi supplemented with the peptide (1 or 10 μM LjRALF34 or LjRALF33) or water (control). While roots were inserted in the liquid, cotyledons were kept outside of the tube. Aluminum foil was used to keep the roots in the dark and to sustain the shoots. After 4 days of incubation (23°C, humidity 50%, 16h/8h light/dark), each plant was extracted and imaged with a digital camera on a light board in a dark chamber.

Regardless of the growing system, the number of lateral roots was manually counted, whereas the length of primary and lateral roots was measured using the Smartroot plugin (Lobét et al. 2011) in ImageJ. Total root length was calculated as the sum of primary and lateral roots length. Lateral root density was calculated by dividing the number of lateral roots for the primary root length.

### Monitoring of Ca^2+^ dynamics with aequorin-based Ca^2+^ reporters

Ca^2+^ measurements in *L. japonicus* roots were performed and analyzed as previously described (Binci et al. 2024). Briefly, after overnight incubation with 5 µM coelenterazine, luminescence from single root segments expressing a cytosol-localized YFP-aequorin-based Ca^2+^ reporter was collected by a custom-built luminometer and converted into Ca^2+^ concentration values. The following stimuli were used: 10^-5^ M or 10^-6^ M synthetic peptide LjRALF34 or LjRALF33, 10^-7^ M CO4 (IsoSep, Tullinge, Sweden), 10^-7^ M CO7 (Elicityl, Crolles, France). All stimuli were dissolved in H_2_O.

### Heterologous expression of LjMLO4, Ca^2+^ measurements, viability assays and imaging in *E. coli* cells

Ca^2+^ assays in *E. coli* were performed as previously described (Teardo et al. 2019). Briefly, chemo-competent C41 *E. coli* were co-transformed via heat-shock with a version of pRham (empty vector, LjMLO4-HIS, LjMLO4Δ-HIS). and the aequorin-expressing pACYCDuet-1(HA1-Aeq) and selected with 50 µg/mL kanamycin and 40 µg/mL chloramphenicol. Liquid cultures were grown at 37°C until OD_600_ = 0.4, when protein expression was induced with 1 mM IPTG (for aequorin), 0.2% (w/v) rhamnose (for LjMLO4s) and 0.25 mM EGTA for 2 h. Bacteria were then resuspended in buffer A (25 mM HEPES, 125 mM NaCl, 1 mM MgCl_2_, 500 μM EGTA, pH 7.5), reconstituted with 5 μM coelenterazine (90 min, room temperature), placed in the luminometer and challenged with Ca^2+^ pulses (1 mM CaCl_2_ in buffer A) or control solution (buffer A only). For LaCl_3_ pre-treatments, root segments were incubated for 10 min with 3 mM LaCl_3_ before the injection of 1 mM CaCl_2_. Luminescence data were analyzed as described for *L. japonicus* Ca^2+^ measurements.

Bacterial cell viability was monitored by the LIVE/DEAD *Bac*Light bacterial viability kit (Thermo Fischer). Transformed bacteria were grown and induced as described for Ca^2+^ assays. Staining was performed according to manufacturers’s protocol with a mixture of SYTO 9 and propidium iodide to distinguish live and dead bacteria. As a positive control (100% dead bacteria), bacteria were incubated for 10 min at 100°C. TEM observations of *E. coli* cells were conducted as described above for *L. japonicus* roots.

Expression of LjMLO4-HIS and LjMLO4Δ-HIS was validated via Western blotting. Bacterial cultures were induced by IPTG and rhamnose for 2 h, 4 h and 16 h and 2 ml aliquots were collected at each timepoint. Cells were pelletted, resuspended in 4x Laemmli Sample Buffer at the same density (100 μl per 1 OD_600_) and incubated at 100 °C for 10 minutes. 20 μl of whole cell lysates were run in a SurePAGE^TM^ Bis-Tris, 4-20% polyacrilamide gel (GenScript, Nanjing, China) and blotted on 0.2 µm nitrocellulose membrane, which was saturated with 5% milk (w/v) in T-TBS (20 mM Tris, 150 mM NaCl, pH 7.5, 0.05% v/v Tween-20). 6xHis-tagged proteins were detected by overnight incubation at 4 °C with HRP-conjugated mouse Anti-His-Tag antibody (SB194b; SouthernBiotech, Birmingham, USA), at 1:20000 dilution in 5% milk T-TBS. Chemiluminescent detection was performed by SuperSignal^TM^ West Pico PLUS developing solution (Thermo Fisher).

The subcellular localization of LjMLO4-HIS and LjMLO4Δ-HIS was validated via immunofluorescence using an anti-His-Tag antibody (Merck), 1:150 diluted, followed by Alexa Fluor 488 goat anti-mouse IgG (Thermo Fisher), as previously described (Zonin et al. 2011).

### Statistical analysis

Data were statistically analyzed and presented graphically using R statistical environment and Rstudio (RStudio Team, 2020; R Core Team, 2022). When passing their assumptions, Student’s *t*-test or the Anova test and Tukey’s post-hoc test were applied; in the other cases, Wilcoxon-Mann-Whitney test or Kruskal Wallis and Dunn’s tests were performed. The statistical tests used are explicit in each figure caption. Only p-values < 0.1 are reported in each figure. Complete raw data and statistical analyses for each figure is reported in Supplementary Table S5.

## Acknowledgements

We thank Andrea Genre (Torino, Italy), Caroline Gutjahr (Potsdam, Germany), Vanessa Checchetto and Ildiko Szabo (Padova, Italy) for fruitful discussions. The technical assistance of staff at the Imaging Facility and the Plant Genome Editing and Phenotyping Facility of the Department of Biology (University of Padova, Italy) is gratefully acknowledged.

## Conflict of interest

The authors declare no conflicts of interest

## Author contributions

MG, LN, FB, conceived the study and designed research; FB and MG performed phylogenetic analysis of MLOs; FB, GR, EDN and GG conducted molecular cloning; FB, EDN and GR genotyped *LORE1* insertional mutants and performed the GUS staining; FB, GG and EDN performed mycorrhization experiments; FB, LN and BB conducted TEM analyses; AC performed phosphate quantification; FB and GG performed root phenotyping; FB, GG and SCS identified RALF peptides in *L. japonicus* and performed treatment experiments with LjRALFs; FB, FV and LC carried out heterologous expression in *E. coli*; FB and GG conducted aequorin-based Ca^2+^ measurements; FB and GG analyzed data, conducted statistical analyses and data visualization; FB prepared figures; FB, MG and LN wrote the text with inputs from GG. All authors read and approved the final manuscript.

## Funding

This work was supported by grants from the European Union (NextGenerationEU 2021 STARS Grants@Unipd program P-NICHE and Next Generation EU, Mission 4, Component 1, PRIN-PNRR 2022 - grant number P2022WL8TS to MG and PRIN 2022 - grant number 2022NW97JX to LN), from the University of Padova (Progetti di Ricerca Dipartimentali - PRID) [grant number BIRD214519 to MG; SEED Giovani 2020 to FB].

## Supplementary data

Supplementary Figure S1. Multisequence alignment of LjMLO protein sequences and predicted structure of LjMLO4.

Supplementary Figure S2. Regulation of p*MLO4* by two different cis-regulator elements (CRE) shown by the GUS assay.

Supplementary Figure S3. Activity of pMLO4 mutated variants in the wild-type genetic background of non-colonized *L. japonicus* roots

Supplementary Figure S4. Validation of the insertional LORE1 mutant lines for the MLO4 locus cDNA extracted from mycorrhizal roots of *L. japonicus*.

Supplementary Figure S5. Root colonization at 7 wpi in wild-type and *mlo4-1* mutant.

Supplementary Figure S6. Representative images of *L. japonicus* wild-type, *mlo4-1* and *mlo4-3* seedlings after treatment with IAA.

Supplementary Figure S7. LjRALFs expression atlas and Interpro predictions.

Supplementary Figure S8. Effect of LjRALFs treatment on primary root length and growth.

Supplementary Figure S9. Controls for Ca^2+^ assays in *E. coli* cells expressing LjMLO4 or LjMLO4Δ.

Supplementary Figure S10. Ca^2+^ measurements in response to LjRALF34.

Supplementary Figure S11. Ca^2+^ measurements in response to LjRALF33.

Supplementary Figure S12. Ca^2+^ measurements in response to abiotic and biotic factors.

Supplementary Table S1. MLO and RALF protein sequences used in this study.

Supplementary Table S2. Sequences of LjRALF mature peptides and LjMLO4 optimize coding sequence used for synthesis.

Supplementary Table S3. List of primers used in this study.

Supplementary Table S4. List of plasmids used in this study.

Supplementary Table S5. Raw data and statistical analysis for each figure

## Data availability

The datasets used in this study are available in the supplementary data.

## Conflict of interest

The authors declare that they have no conflicts of interest.

## Notes

### Competing Interest Statement

The authors have declared no competing interest.

### Summary of Updates

The manuscript has been updated to increase robustness of the findings that LjMLO4 functions both in root development and in arbuscular mycorrhiza. We added gene expression analyses of LjMLO4 and AM marker genes in mycorrhizal roots (Fig 1 and 2), dose-dependent effect of auxin on the mlo4 mutant root phenotypes (Fig.3), normalized analysis of root growth in response to RALF peptides (Fig. 4). Pharmacological, viability and imaging controls further corroborated calcium bioassays in E. coli cells showing that LjMLO4 mobilizes calcium across membranes (Fig. 5 and S9).

