## Supplementary Figures for "A symbiotic MLO gene regulates root development via RALF34-triggered Ca^2+^ signalling in *Lotus japonicus*"

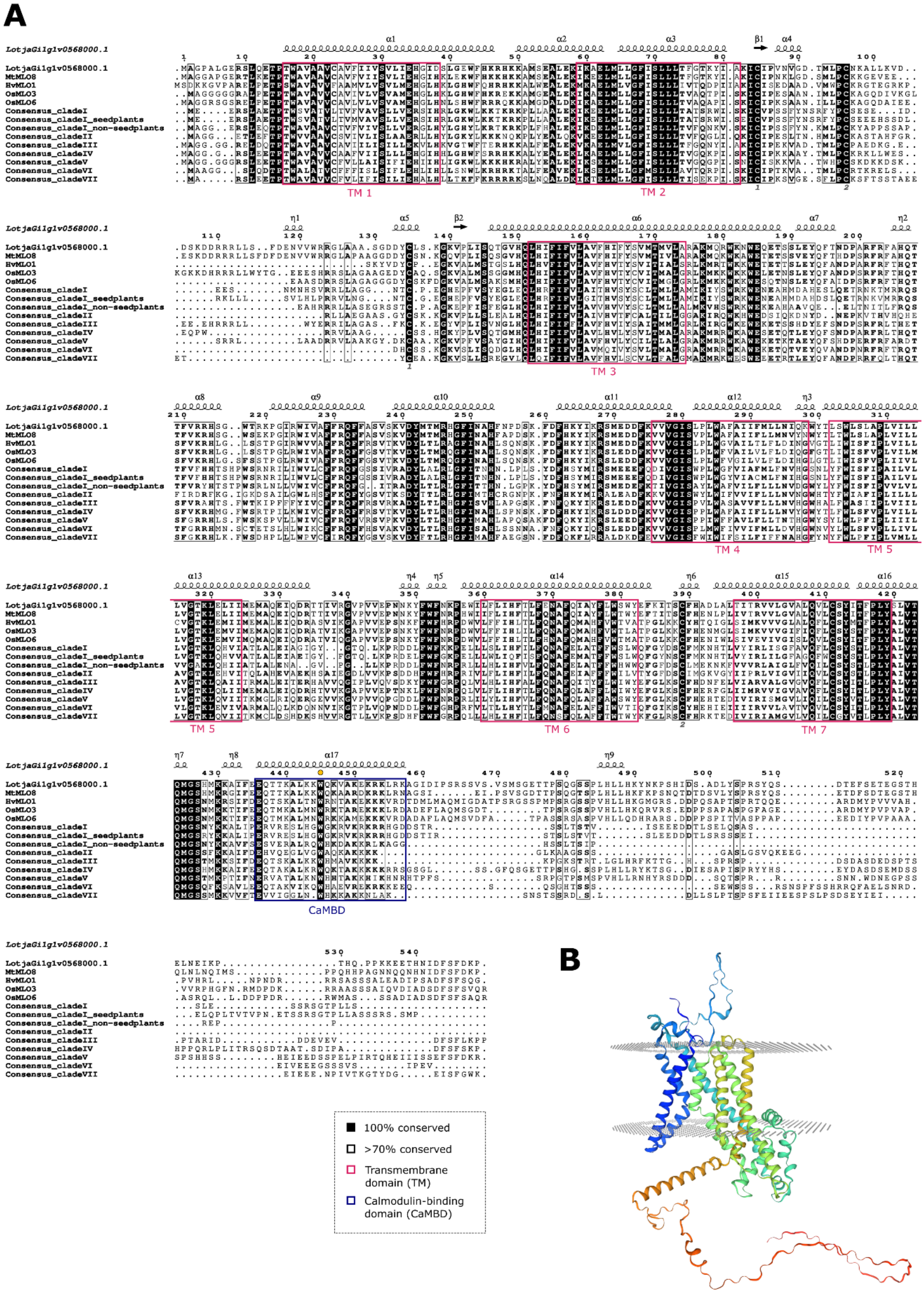
A
LotjaGi1g1v0568000.1
MtMLO8
HvMLO1
OsMLO3
OsMLO6
Consensus_cladeI
Consensus_cladeI_seedplants
Consensus_cladeI_non-seedplants
Consensus_cladeII
Consensus_cladeIII
Consensus_cladeIV
Consensus_cladeV
Consensus_cladeVI
Consensus_cladeVII
TM 1
TM 2
TM 3
TM 4
TM 5
TM 6
TM 7
CaMBD
B
100% conserved
>70% conserved
Transmembrane domain (TM)
Calmodulin-binding domain (CaMBD)

**Supplementary Figure S1. Multisequence alignment of LjMLO protein sequences and predicted structure of LjMLO4.** A) Alignment with the ClustalOmega algorithm of the consensus protein sequence of clades I, II, III, IV, V, VI, VII MLOs (retrieved from Kusch et al. 2016) with clade IV MLO proteins from *L. japonicus, M. truncatula, H. vulgare and O. sativa*. Conserved residues are indicated by solid black (100% conserved) and black-outlined (>70% conserved) boxes. The 7 transmembrane domains (TM) are highlighted in magenta, and the calmodulin-binding domain (CaMBD) is shown in blue. The conserved tryptophan residue within the CaMBD is marked by a yellow dot, and the two pairs of conserved cysteine residues predicted to form disulfide bonds are indicated by matching numbers at the bottom of the alignment. The predicted secondary structure of LjMLO4 is shown on top (α, α-helices; β, β-strands; η, $3_{10}$-helices). Multisequence alignment was visualized using ESPript. B) Structure prediction of LjMLO4 computed via the Swiss-Model server. The model includes a schematic representation of a cell membrane bilayer (grey; top, outer leaflet; bottom, inner leaflet). The colour gradient of the protein model is arbitrary.

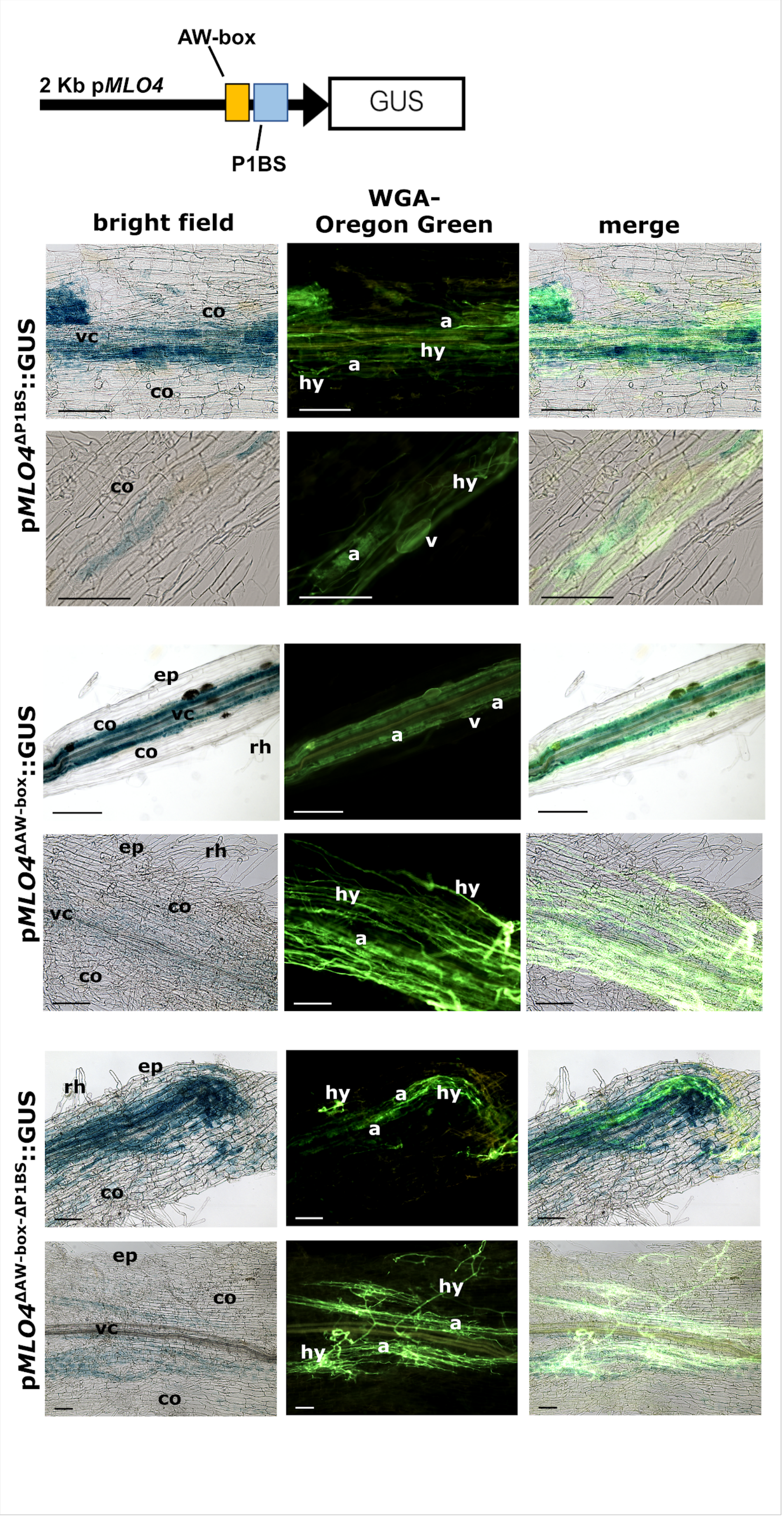
AW-box
2 Kb pMLO4
GUS
P1BS
bright field
WGA-Oregon Green
merge
pMLO4ΔP1BS::GUS
pMLO4ΔAW-box::GUS
pMLO4ΔAW-box-ΔP1BS::GUS
vc
co
a
hy
v
ep
rh

**Supplementary Figure S2. Regulation of p*MLO4* by two different *cis*-regulator elements (CRE) shown by the GUS assay.** The CREs AW-box and P1BS found in p*MLO4* (scheme at the top left) were mutated either alternatively (p*MLO4*$^{\Delta P1BS}$ or p*MLO4*$^{\Delta AW\text{-}box}$) or jointly (p*MLO4*$^{\Delta AW\text{-}box\text{-}\Delta P1BS}$). Transgenic composite *L. japonicus* plants were generated expressing the *UidA* gene under the control of the different versions of the promoter. Plants were co-cultivated for 5 weeks with *R. irregularis* at a low phosphate regime. Harvested fresh roots were stained with GUS solution overnight, fixed in 70% ethanol, clarified in 10% KOH and counter-stained with WGA-Oregon Green. Representative images from multiple root samples are shown. a, arbuscule; co, cortex; ep, epidermis; hy, hypha; rh, root hair; v, vesicle; vc, vascular cylinder. Bar, 100 µm.

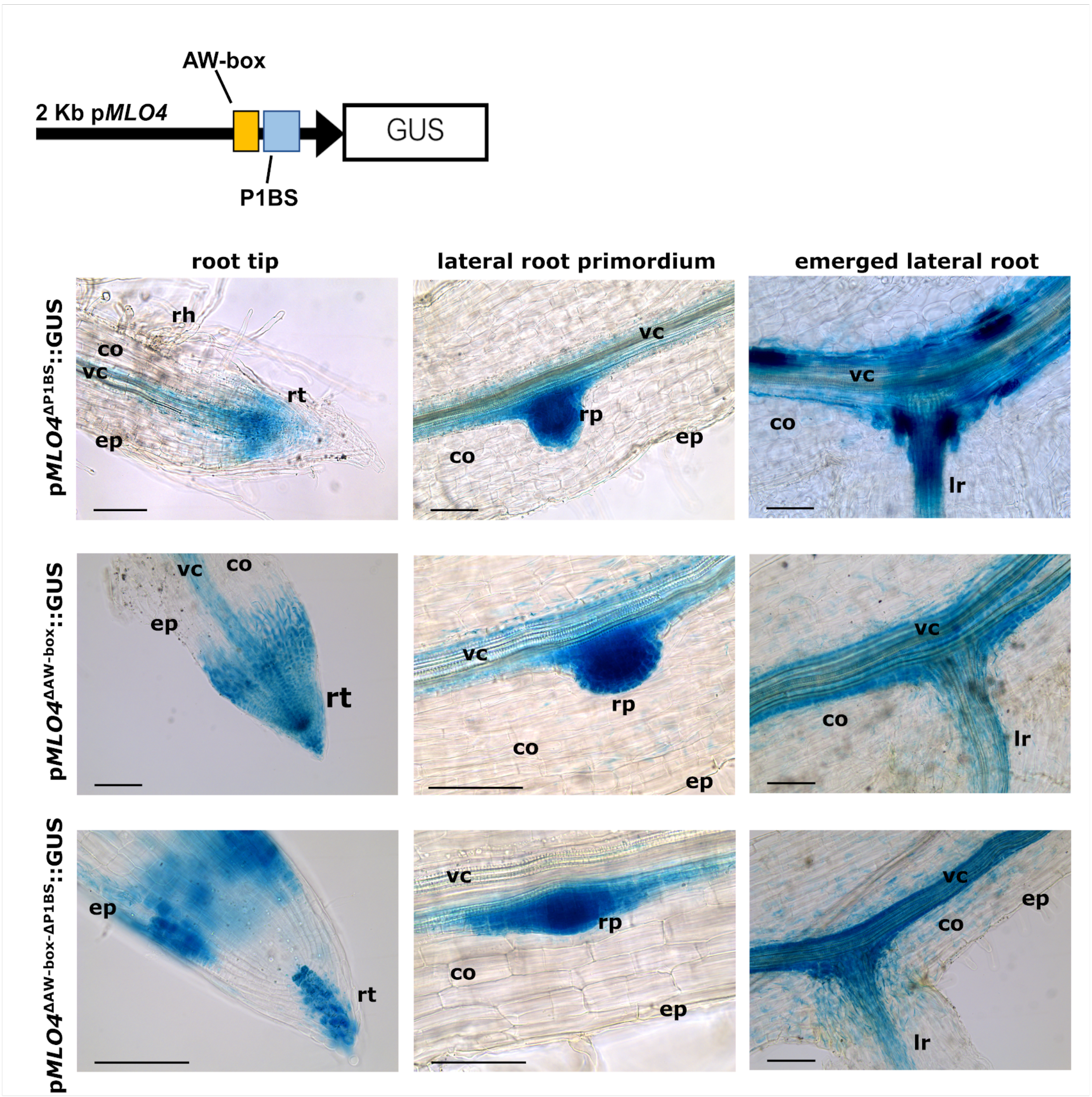


**Supplementary Figure S3. Activity of p*MLO4* mutated variants in the wild-type genetic background of non-colonized *L. japonicus* roots.** Composite plants generated with *A. rhizogenes*-mediated hairy roots transformation were grown in pots for 5 weeks in the absence of AM fungi. Fresh roots were stained with GUS solution, fixed in 70% ethanol and clarified in 10% KOH before imaging. co, cortex; ep, epidermis; lr, lateral root; rh, root hair; rp, root primordium; rt, root tip; vc, vascular cylinder. Bar, 100 µm.

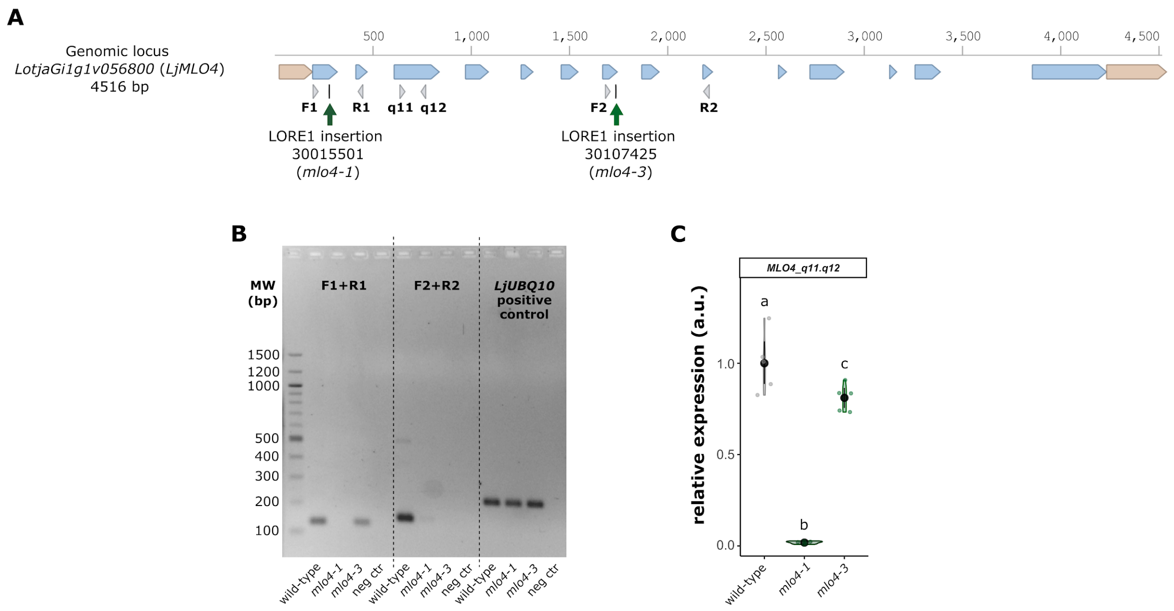


**Supplementary Figure S4. Validation of the insertional LORE1 mutant lines for the *MLO4* locus on cDNA extracted from mycorrhizal roots of *L. japonicus*.** A) The schematic representation of the genomic locus *MLO4* is shown with 5' and 3' UTRs (brown), exons (blue), introns (blank spaces between exons), sites of LORE1 insertions (black lines) and the primers used for genotyping (grey arrowheads). B) The results of semi-quantitative RT-PCR amplifications (30 cycles) from wild-type, *mlo4-1* and *mlo4-3* cDNA after electrophoretic run in 1.5% agarose gels are shown. The *LjUbiquitin10* housekeeping gene (*LjUBQ10*) was used as a positive control. In each reaction, water was used as a negative control (neg ctr). C) Quantification of *LjMLO4* via RT-qPCR performed on the same cDNA. Different letters indicate significant differences among genotypes.

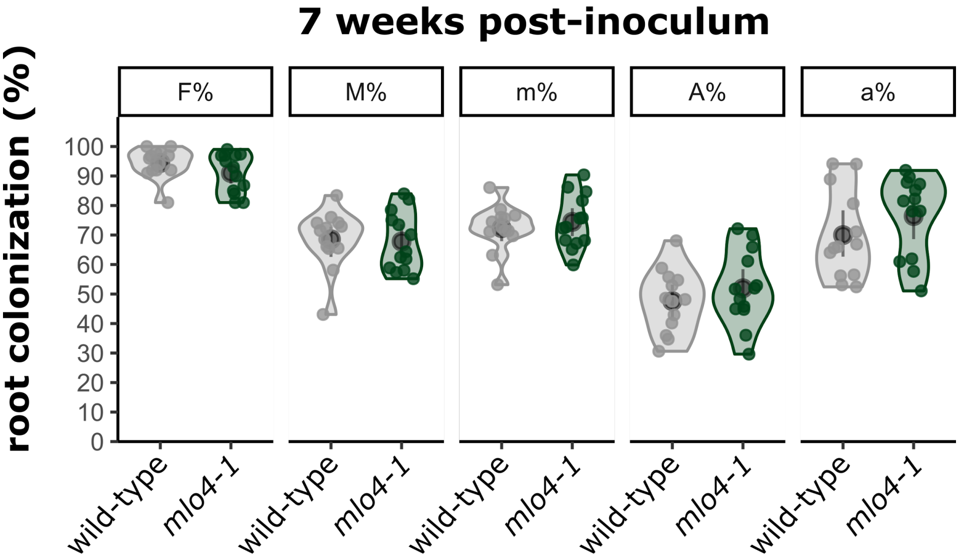


**Supplementary Figure S5. Root colonization at 7 wpi in wild-type and *mlo4-1* mutant.** Quantification of fungal colonization in wild-type, *mlo4-1 L. japonicus* roots after 7 weeks of co-cultivation with *R. irregularis.* Colonization parameters were calculated according to the Trouvelot method: F%, frequency of mycorrhiza; M%, absolute intensity of mycorrhiza; m%, relative intensity of mycorrhiza; A%, arbuscule abundance; a%, relative arbuscule abundance. The distribution of data is presented as violin plots, each dot is a biological replicate. No significant differences were identified after ANOVA.

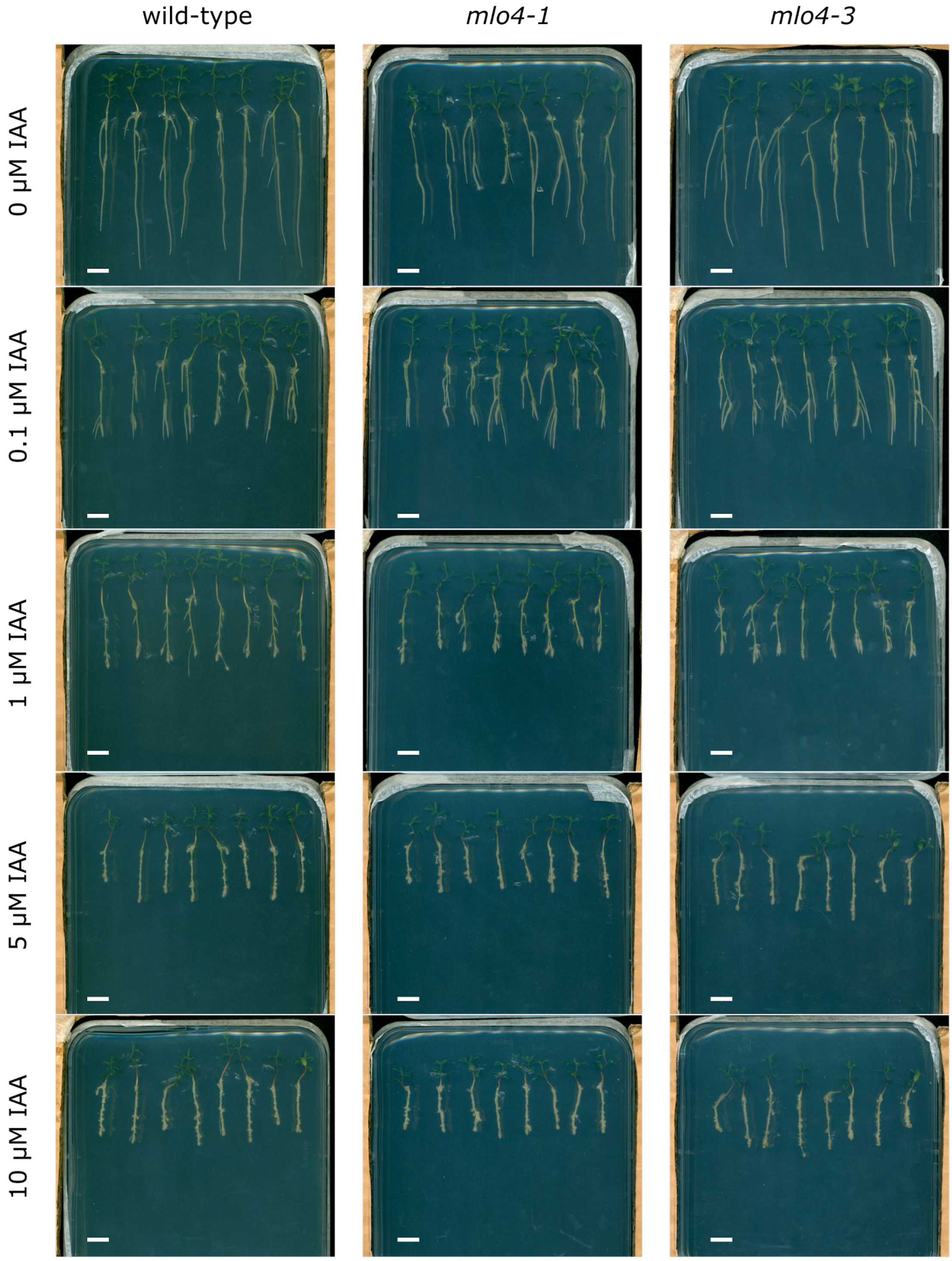


**Supplementary Figure S6. Representative images of *L. japonicus* wild-type, *mlo4-1* and *mlo4-3* seedlings after treatment with IAA.** Representative images (N = 40 seedlings per genotype and condition) of wild-type, *mlo4-1* and *mlo4-3* seedlings after 10 days of treatment with 0, 0.1, 1, 5, 10 µM IAA. Bar, 1 cm.

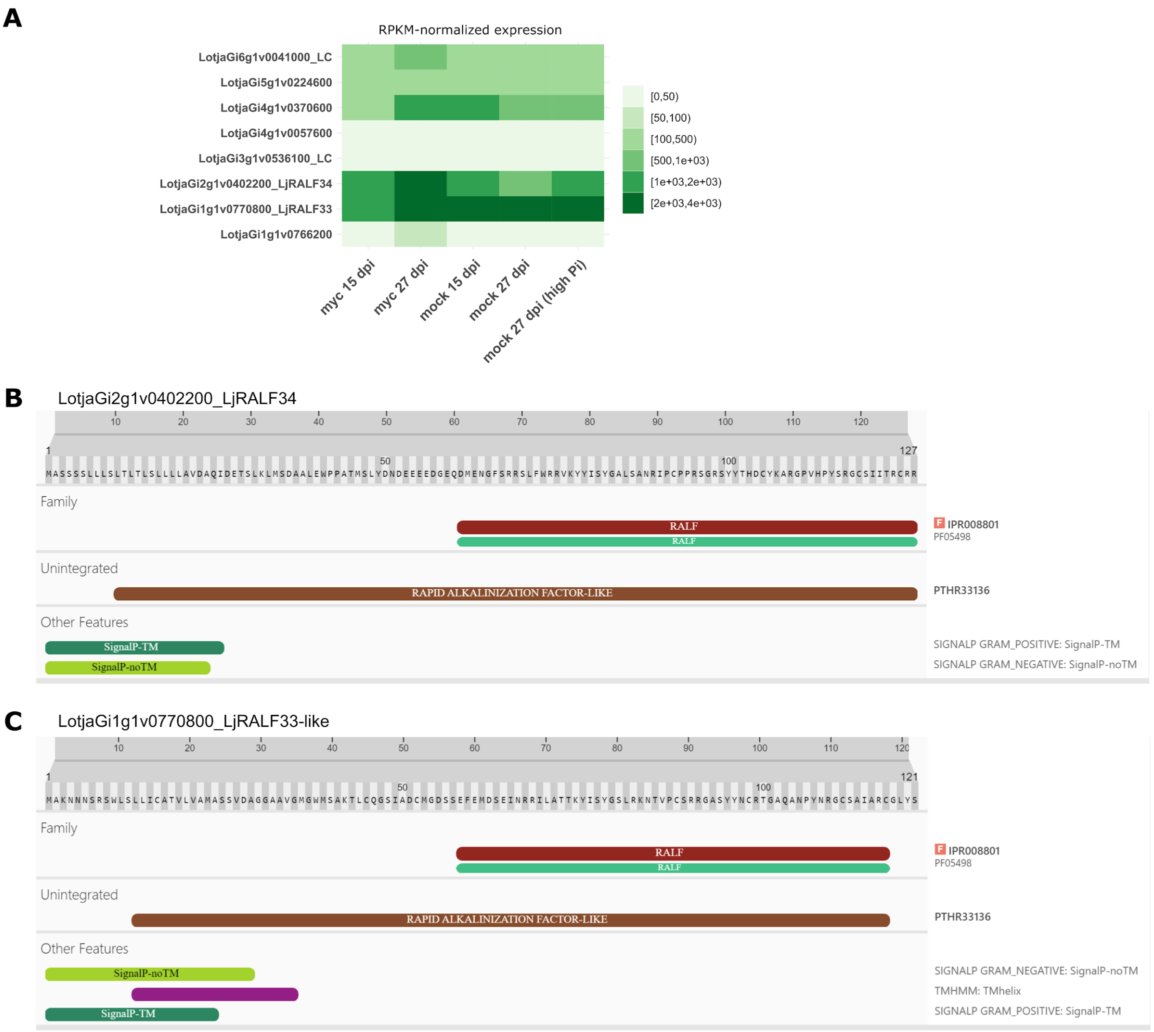


**Supplementary Figure S7. LjRALFs expression atlas and Interpro predictions.** A) Raw expression data as heatmap were downloaded from *L. japonicus* ExpressionAtlas in LotusBase, using the Gifu v1.2 genome dataset (Kamal et al. 2020). Rows represent the 8 *LjRALF*-like genes and columns are the different conditions from the dataset considered. Different shades of green represent the expression level (raw data) in the different considered conditions (darker green means higher transcript abundance). B-C) Interpro prediction of functional domains in the full-length protein sequences of LjRALF34 (B) and LjRALF33 (C).

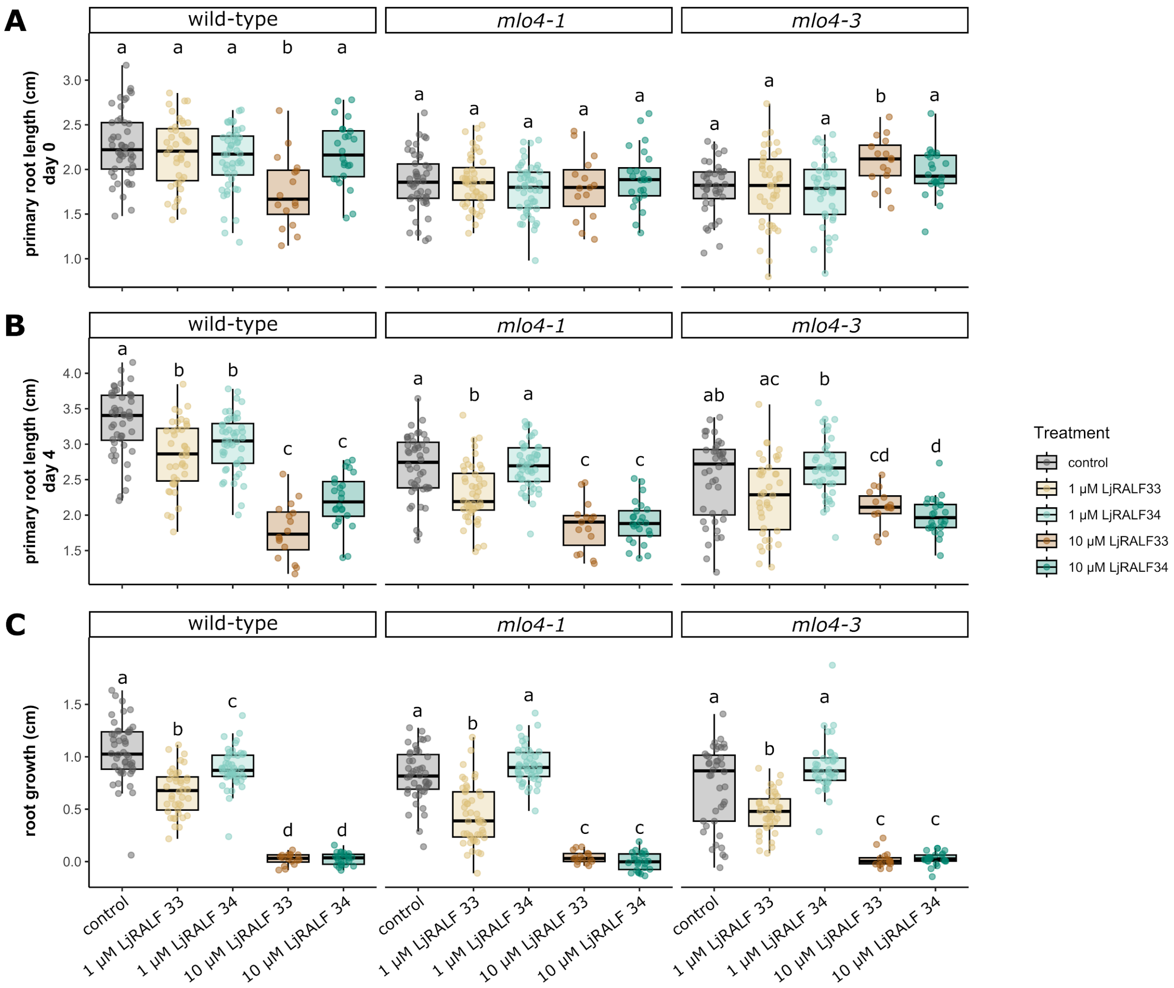


**Supplementary Figure S8. Effect of LjRALFs treatment on primary root length and growth.** 7-day-old *L. japonicus* seedlings of different genotypic backgrounds (wild-type*, mlo4-1 and mlo4-3* lines) were treated with two different concentrations (1 µM and 10 µM) of LjRALF34 and LjRALF33 or grown in control conditions (liquid modified LA medium with 200 µM $P_i$). Each plant was imaged before and after 4 days of treatment and the primary root length was quantified with the ImageJ plugin Smartroot. Length before (day 0, A) and after (day 4, B) the treatment and the difference between the two (root growth, C). Each dot represents a biological replicate and the data distribution as violin plots. Statistical analysis was performed using Kruskal-Wallis, followed by Dunn's post-hoc for pairwise comparisons. Different letters indicate significantly different statistical groups within each treatment.

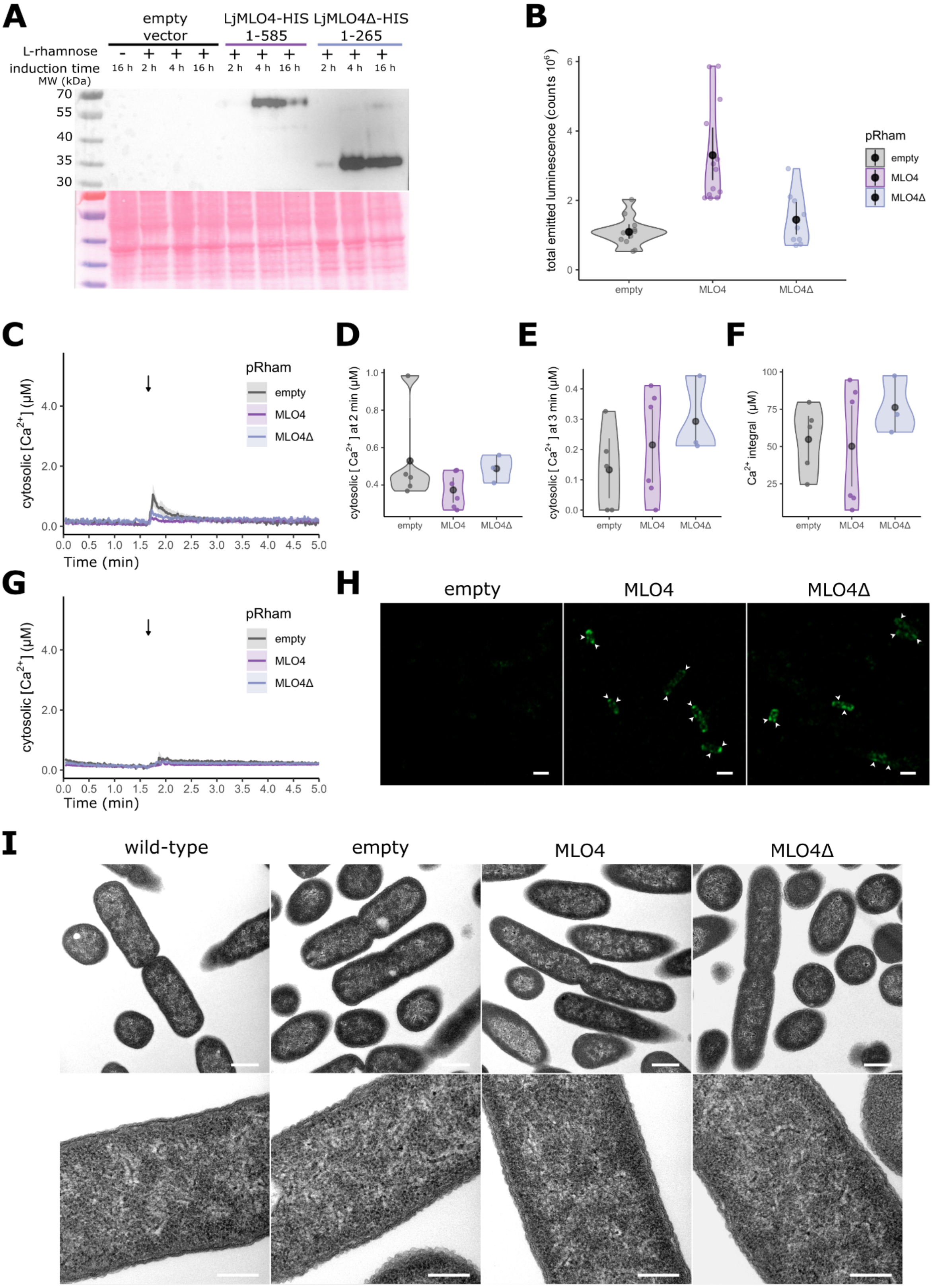

A
empty vector
LjMLO4-HIS 1-585
LjMLO4Δ-HIS 1-265
L-rhamnose
induction time
MW (kDa)
- + + + + + + + + +
16 h 2 h 4 h 16 h 2 h 4 h 16 h 2 h 4 h 16 h
70
55
40
35
30
B
total emitted luminescence (counts 10^6)
pRham
empty
MLO4
MLO4Δ
C
cytosolic [Ca2+] (μM)
Time (min)
D
cytosolic [Ca2+] at 2 min (μM)
E
cytosolic [Ca2+] at 3 min (μM)
F
Ca2+ integral (μM)
G
H
empty
MLO4
MLO4Δ
I
wild-type
empty
MLO4
MLO4Δ

**Supplementary Figure S9. Control experiments for $Ca^{2+}$ assays in *E. coli* cells expressing LjMLO4 or LjMLO4Δ.** A) Western blot showing the expression of LjMLO4-HIS and LjMLO4Δ-HIS in *E. coli* cells using an anti-HIS antibody. Equal loading was confirmed by Ponceau Red staining of the blot membrane (bottom). B) Total emitted luminescence from the bacterial samples (50 µl) expressing the three different pRham expression constructs. C-F) $Ca^{2+}$ changes after injection of control solution (buffer A only) represented as $Ca^{2+}$ traces along time (C), concentration of $Ca^{2+}$ after 2 min (D) and after 3 min (E), total $Ca^{2+}$ mobilized over 5 minutes (F). No statistical differences were observed after Kruskal-Wallis test followed by Dunn's post-hoc correction. G) $Ca^{2+}$ traces in response to control solution (buffer A only) after a 10 min-long pre-treatment with 3 mM$LaCl_3$; data are presented as means ± SE (shading). H) Immunofluorescence localization of LjMLO4-HIS and LjMLO4Δ-HIS in bacterial cells using anti-HIS (primary antibody) and anti-mouse IgG-AlexaFluor488 (secondary antibody). Bar, 2 µm. I) Transmission Electron Microscopy (TEM) observations of *E. coli* cells (wild-type and expressing the three different pRham expression constructs), showing good preservation of bacterial ultrastructure. Bar, 500 nm (top) and 200 nm (bottom).

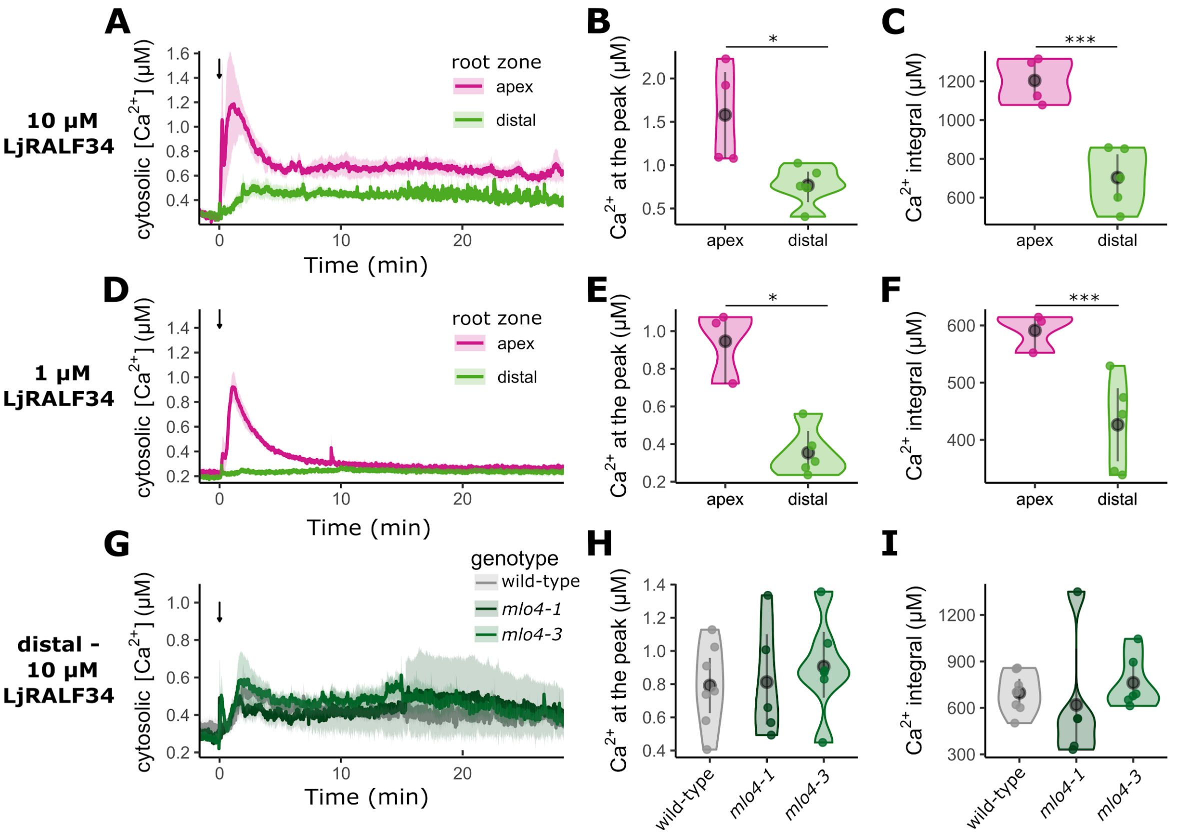


**Supplementary Figure S10. $Ca^{2+}$ measurements in response to LjRALF34.** $Ca^{2+}$ assays were performed with *L. japonicus* root segments expressing the cytosolic $Ca^{2+}$ reporter YFP-aequorin after stimulation with 10 and 1 µM LjRALF34. A-F) $Ca^{2+}$ assays were conducted in *L. japonicus* wild-type root segments expressing a cytosolic targeted aequorin probe and including the root apex (pink) or excised more distal from it (light green). G-H) $Ca^{2+}$ measurements in response to 10 µM LjRALF34 in root segments distal from the root apex. Comparison among wild-type, *mlo4-1* and *mlo4-3* genetic backgrounds are shown. A, D, G) data are presented as means ± SE (shading) of $n \geq 3$ obtained from at least 3 different composite plants (independent transformations). The stimulus was injected at time 0 (min), indicated by the black arrow. B, E, H) dots represent the maximum $[Ca^{2+}]$ for each run. C, F, I) dots represent the total mobilized $[Ca^{2+}]$ ($Ca^{2+}$ integral) for each run (30 min). Statistical analysis was conducted via Wilcoxon-Mann-Whitney test (B, E), t-test (C, F) or Kruskal-Wallis non-parametric test followed by Dunn's post-hoc correction for pairwise comparisons (H, I). Only p-values <0.1 are reported, *$p<0.05$; *** $p<0.001$.

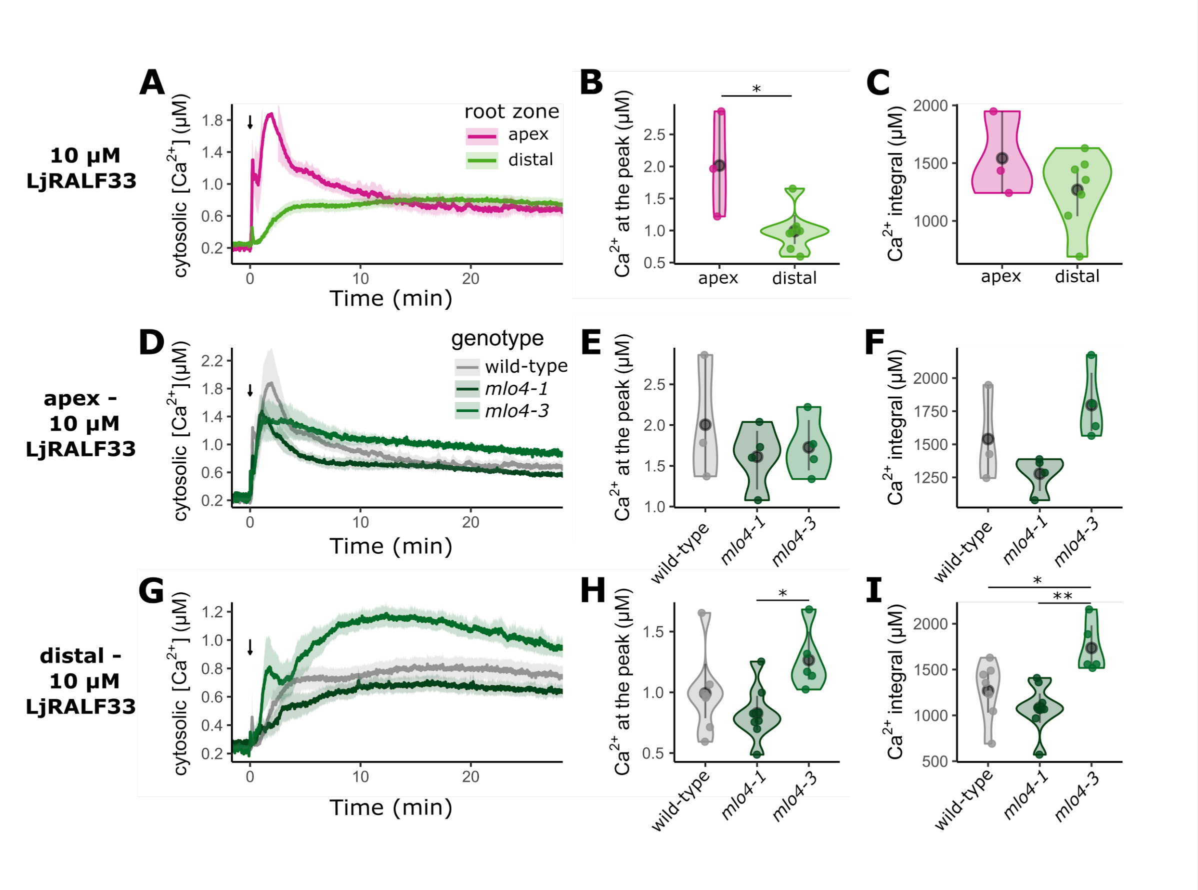


**Supplementary Figure S11. $Ca^{2+}$ measurements in response to LjRALF33.** $Ca^{2+}$ assays were performed with *L. japonicus* root segments expressing the cytosolic $Ca^{2+}$ reporter YFP-aequorin after stimulation with 10 μM LjRALF33. A-C) $Ca^{2+}$ measurements conducted in root segments including the root apex (pink) or excised more distal from it (light green). D-I) Comparison among wild-type, *mlo4-1* and *mlo4-3* genetic backgrounds are shown. In A, D, G), [$Ca^{2+}$] changes over time are presented as means ± SE (shading) of n≥3 traces from at least three different composite plants (independent transformations). The stimulus was injected at time 0 (min), indicated by the black arrow. In B, E, H), dots represent the maximum [$Ca^{2+}$] for each trace in the whole run. In C, F, I), dots represent the total mobilized [$Ca^{2+}$] ($Ca^{2+}$ integral) during the whole run (30 min). Statistical analysis was conducted via Wilcoxon-Mann-Whitney test (B, E), t-Test (C, F) or Kruskal-Wallis non-parametric test followed by Dunn's post-hoc correction for pairwise comparisons (H, I). *p<0.05; **p<0.01.

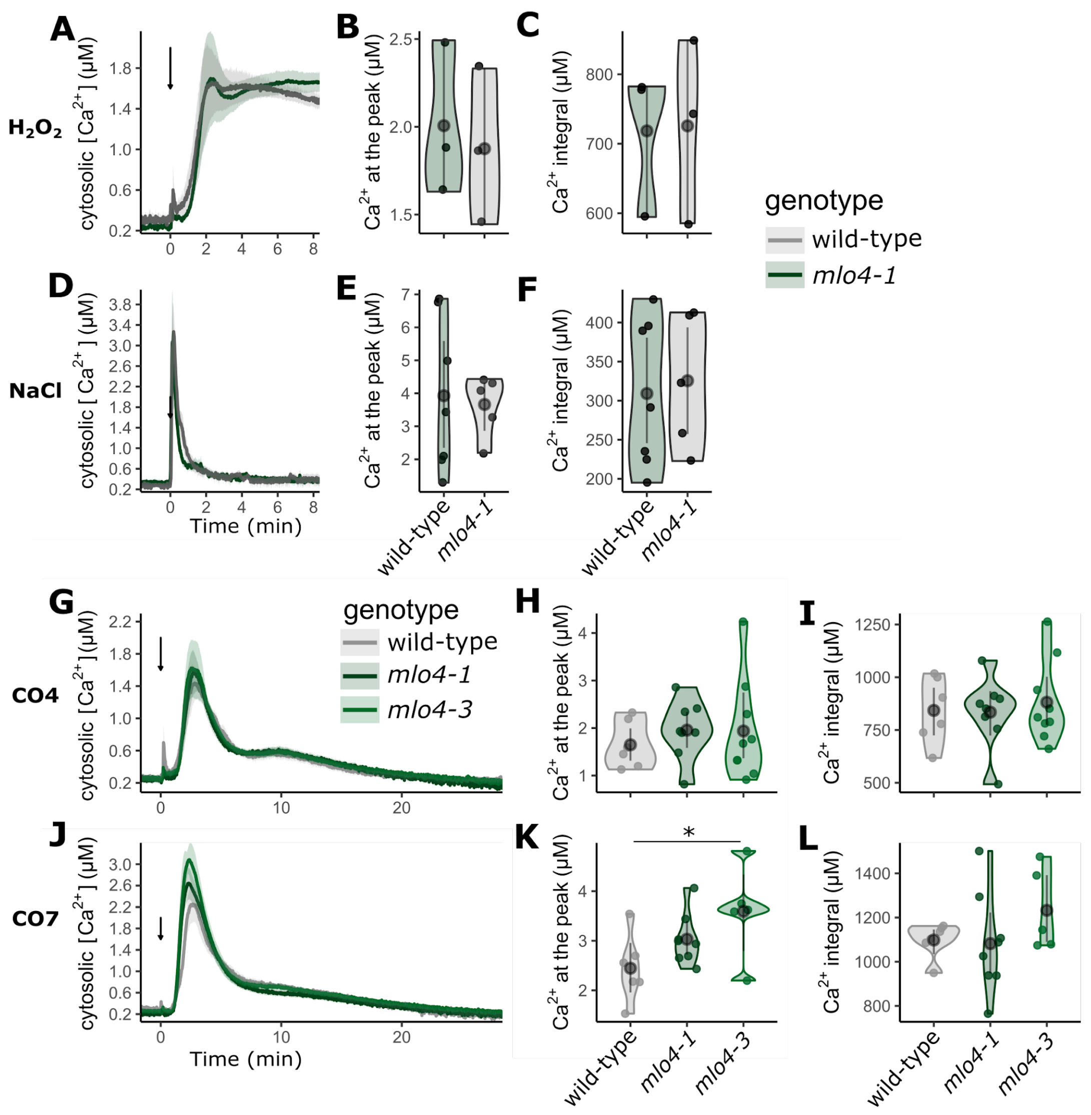


**Supplementary Figure S12. $Ca^{2+}$ measurements in response to abiotic and biotic factors.** $Ca^{2+}$ assays were performed in wild-type, *mlo4-1, mlo4-3 L. japonicus* root segments expressing the cytosolic $Ca^{2+}$ reporter YFP-aequorin after stimulation with 10 mM $H_2O_2$ (A-C), 0.1 mM NaCl (D-F), $10^{-7}$M CO4 (G-I), $10^{-7}$M CO7 (J-L). In A, D, G, J, [$Ca^{2+}$] changes over time are presented as means ±SE (shading) of n≥3 traces from at least three different composite plants (independent transformations). The stimulus was injected at time 0 (min), indicated by the black arrow. In B, E, H, K, dots represent the maximum [$Ca^{2+}$] for each trace in the whole run. In C, F, I, dots represent the total mobilized [$Ca^{2+}$] ($Ca^{2+}$ integral) during the whole run. Statistical analysis was conducted via *t*-test (B, C, E, F) or ANOVA test (H, I, K, L) followed by Tukey's post-hoc correction. $*p<0.05$.
