## Supplementary tables S2-S3-S4 for "A symbiotic MLO gene regulates root development via RALF34-triggered Ca^2+^ signalling in *Lotus japonicus*"

**Supplementary Table S2. Sequences of LjRALF mature peptides and *LjMLO4* optimized coding sequence used for synthesis.**

| | |
|---|---|
| LjRALF34 | FWRRVKYYISYGALSANRIPCPPRSGRSYYTHDCYKARGPVHPYSRGCSIITR<br>CRR |
| LjRALF33 | ATTKYISYGSLRKNTVPCSRRGASYYNCRTGAQANPYNRGCSAIARCGLYS |
| *LjMLO4* codon optimized coding sequence fused to HIS-tag and STREP-tag at the 3' end | ATGGCCGGTCCTGCATTAGGCGAGCGTTCATTACAGGAAACCCCTACCTG<br>GGCGGTGGCAGCAGTTTGCGCCGTTTTCATAATTGTGTCAGTACTGATAG<br>AGCACGGCATTGACTCGCTGGGTGAATGGTTCCATAAGCGTCACAAGAA<br>GGCAATGTCAGAGGCACTCGAAAAGATAAAGGCCGAACTGATGTTGCTG<br>GGCTTTATCTCTTTACTCCTGACCTTTGGAACGAAGTACATCGCGAAAAT<br>TTGTATACCGGTCAACGTGGGCGACACAATGCTGCCCTGTAACAAAGCTC<br>TCTTAAAGGTGGACGATAGTAAGGACGACCGCCGGCGACTCTTAAGTTTT<br>GACGAGAACGTCGTTTGGCGAAGAGGATTGGCCGCCGCTAGCGGCGACG<br>ATTACTGCCTGTCTAAGGGTAAGGTACCGCTTATTTCCCAGACGGGCGTA<br>CATCAGCTTCACATCTTCATCTTTGTATTAGCTGTGTTTCATATCTTTTATT<br>CGGTAATGACAATGGTACTTGCTCGCGCGAAGATGCAACGCTGGAAGAA<br>CTGGGAGCAGGAGACGTCGAGCCTCGAATACCAGTTCACGAACGACCCG<br>GCGCGATTTCGGTTCGCACATCAGACCACTTTTGTGCGACGCCATAGCGG<br>TTGGACCCGTAAGCCGGGGATCCGCTGGATCGTCGCTTTCTTTCGTCAGT<br>TCTTCGCATCAGTATCAAAAGTTGACTATATGACGATGCGTCACGGTTTC<br>ATCAATGCGCACTTCAACCCGGACTCCAAGTTCGATTTCCACAAGTATAT<br>TAAGCGCAGCATGGAAGACGACTTCAAGGTAGTTGTGGGCATTTCATTAC<br>CCCTCTGGGCATTCGCTATTATATTTATGCTCTTAAATATCCAAAAGTGGT<br>ATACCCTTTCATGGTTGAGTCTTGCTCCGCTGGTTATTCTGCTGTTGGTTG<br>GCACAAAATTGGAGCTGATTATAATGGAGATGGCGCAGGAGATTCAGGA<br>CCGCACCACCATAGTCCGTGGTGTGCCTGTTGTGGAACCTAATAATAAAT<br>ACTTTTGGTTCAACAAACCCGAGTGGATACTGTTTCTGATTCACTTCACTC<br>TCTTCGAAAACGCTTTTCAGATCGCCTACTTTCTCTGGTCTTGGTACGAAT<br>TTAAAATTACCAGTTGTTTTCACGCGGATTTAGCCTTGACTATCACCCGCG<br>TGGTTCTGGGAGTTGCGCTGCAGGTGCTGTGTAGCTATATAACCTTCCCA<br>CTGTACAGTCTGGTTACCCAAATGGGTAGCCATATGAAGAAGGCGATTTT<br>CGAAGAGCAGACTACTAAAGCCCTGAAGAAATGGCAGAAAGTAGCTAAA<br>GAGAAACGAAAACTGCGAAAGGCGGGCATCGACATACCGAGTCGTTCTT<br>CGGTATCGGTATCTATGTCTGGTGAAACTACGCCGTCTCAGGGTTCTTCTC<br>CCCTGCACCTGTTGCATAAATATAATAAACCGTCCCATATCGATAGCGCG<br>GACCTGTACTCCCCTCGTTCTTACCAGTCAGATACTGAATTCTCTGAGAC<br>GGAAGGTAGTACCCATGAGTTGAACGAGATTAAGCCGACTCACCAGCCG<br>CCGAAGAAAGAGGAGACGCATAACATTGACTTCTCCTTCGATAAACCAG<br>GCTCCCACCACCATCACCATCACTGGAGCCACCCGCAGTTCGAAAAGTAG<br>TAG |

**Supplementary Table S3. List of primers used in this study.**

| Name | 5’ → 3’ sequence | Description |
|---|---|---|
| q81 | CAATGTCGCCAAGGCCCATGGTG | qPCR FOR primer for *LjATPase* (Binci et al. 2024) |
| q82 | AACACCACTCTCGATCATTTCTCTG | qPCR REV primer for *LjATPase* (Binci et al. 2024) |
| q11 | GATTCATATCCCTGTTACTT | qPCR FOR primer for *LjMLO4* |
| q12 | TAACCTCCTCCTATCATCT | qPCR FOR primer for *LjMLO4* |
| PT4F | GTACAATGACCTCATGGTTCT | qPCR FOR primer for *LjPT4* (Volpe et al. 2016) |
| PT4R | CGTTCATCTCGAAATCCTTATC | qPCR REV primer for *LjPT4* (Volpe et al. 2016) |
| q41 | CACGTTGTTAGGACCCCAAT | qPCR FOR primer for *LjSbtM1* (Pimprikar et al. 2015). |
| q42 | TTGAGCAGCACCCTCTCTATC | qPCR REV primer for *LjSbtM1* (Pimprikar et al. 2015) |
| P2 | CCATGGCGGTTCCGTGAATCTTAGG | LORE1 insertion REV |
| G28 | TTCGCTCATGGCCTTCTTATGGCG | *mlo4-1* FOR |
| G29 | AAAGAGGCTTGTTACGGTATCGGCAG | *mlo4-1* REV |
| G67 | TCGTTAGGCGTCACTCAGGCTGGA | *mlo4-3* FOR |
| G81 | TGCTATCAGGATTAAAATGTGCCTGCAT | *mlo4-3* REV |
| Ubi_F | ATGCAGATCTTCGTCAAGACCTT | LjUbiquitin10 FOR |
| Ubi_R | ACCTCCCCTCAGACGAAG | LjUbiquitin10 REV |
| C16 | AACAGGTCTCAACCTGGGTAGAGTTATGTCTA ACTCACTG | modA_MLO4_promoter for |
| C17 | AACAGGTCTCATGTTGTTAATTTTGCTCTCAGC TTCTCC | modA_MLO4_promoter rev |
| C155 | AACAGGTCTCAACCTGTCAAACCGCGCCTAAC ATTT | pMLO4(-2000-208)_modA_for |
| C156 | AACAGGTCTCATGTTTTTACAAGTTAATTTATT GTTTTAGAATTTTTCTTAAACA | pMLO4(-2000-208)_modA_rev |
| C157 | AACAGGTCTCAAACATTATATTAATTAATCAAT TATGTCTTGT | pMLO4(-186-1)_ modB_for |

| | | |
|---|---|---|
| C158 | AACAGGTCTCAAGCCGTTAATTTTGCTCTCAGCTT | pMLO4(-186-1)_ modB_rev |
| C159 | AACAGGTCTCAAACAGCATATGCTTATATTAATTAATCAATTATG | pMLO4(-AW-Box)_ modB_for |
| C160 | AACAGGTCTCATGTTCGGCCTATGCAAGGTTTACAAG | pMLO4(-2000-208+AW-box)_ modB_rev |
| C239 | ACCGCTGAGCAATAAC | pRham linearization FOR |
| C240 | ATGTATATCTCCTTCTTATAGTTAAAC | pRham linearization REV |
| C243 | TATAAGAAGGAGATATACATATGGCCGGTCCTGCATTAGG | LjMLO4_optimised linearization FOR |
| C244 | GCTAGTTATTGCTCAGCGGTCTACTACTTTTCGAACTGCGGGTGGCTC | LjMLO4_optimised linearization REV |
| C285 | TGGTGATGGTGGTGGGAGCCACCGTGACGCATCGTCATATAG | LjMLO4Δ_optimised linearization FOR |

**Supplementary Table S4. List of plasmids used in this study**

| Name | Type | Resistance | SOURCE | Description |
|---|---|---|---|---|
| pGGA000 | entry | ampicillin and chloramphenicol | Addgene - GreenGate cloning system | Empty entry vectors (ccdB+ ) - ModA |
| pGGA006 | entry | ampicillin | Addgene - GreenGate cloning | Arabidopsis UBQ10 promoter - Module A |
| GG59 | entry | ampicillin | Binci *et al.* 2024 | *L. japonicus* 2200bp UBQ10 promoter - Module A |
| GG83 | entry | ampicillin | This study | 2000bp promoter of LjMLO4 - ModA |
| GG148 | entry | ampicillin | This study | pMLO4_modA_-2000_-208 - modA |
| pGGB000 | entry | ampicillin and chloramphenicol | Addgene - GreenGate cloning system | Empty entry vectors (ccdB+ ) - Module B |
| pGGB003 | entry | ampicillin | Addgene - GreenGate cloning | Dummy – module B |
| GG145 | entry | ampicillin | This study | pMLO4_minusAW-box_modB_-1_-224 - ModB |
| GG146 | entry | ampicillin | This study | pMLO4_minusAW-box_minusP1BS_modB_-1_-208 - ModB |
| GG147 | entry | ampicillin | This study | pMLO4_plusAW-box_modA_-2000_-208 - modB |

| | | | | |
|---|---|---|---|---|
| pGGC051 | entry | ampicillin | Addgene - GreenGate cloning system | *UidA* gene for GUS staining - modC |
| pGGD002 | entry | ampicillin | Addgene - GreenGate cloning system | Dummy – module D |
| pGGE009 | entry | ampicillin | Addgene - GreenGate cloning system | Arabidopsis UBQ10 terminator - Module E |
| GG69 | entry | ampicillin | This study | Transformation marker cassette: pAtUBQ10::GFP - modF |
| GG108 | destination | kanamycin | Binci et al. 2024 | Empty destination vector (ccdB+) |
| GG62 | expression | kanamycin | Binci et al. 2024 | cytosolic YFP-linker-aequorin under the control of the LjUBQ10 promoter. Transformation marker: cytosolic mCherry. Destination vector: GG108. |
| GG88 | expression | kanamycin | This study | GUS under p*MLO4.* Transformation marker: cytosolic GFP. Destination vector: GG108. |
| GG142 | expression | kanamycin | This study | GUS under p*MLO4*ΔP1BSΔAW-box. Transformation marker: cytosolic GFP. Destination vector: GG108. |
| GG143 | expression | kanamycin | This study | GUS under p*MLO4*ΔP1BS. Transformation marker: |

| | | | | |
|---|---|---|---|---|
| | | | | cytosolic GFP. Destination vector: GG108. |
| GG144 | expression | kanamycin | This study | GUS under p*MLO4*ΔAW-box. Transformation marker: cytosolic GFP. Destination vector: GG108. |
| pRham | destination | kanamycin | Lucigen (Middleton, WI, USA) | Empty destination vector - pRham |
| pRham-LjMLO4 | expression | kanamycin | This study | LjMLO4-HIS optimized for *E. coli* expression. Rhamnose-inducible expression |
| pRham-LjMLO4Δ | expression | kanamycin | This study | LjMLO4Δ-HIS optimized for *E. coli* expression. Rhamnose-inducible expression |
| pACYC Duet-1(HA1-Aeq) | expression | chloramphenicol | Teardo et al. 2019 | Aequorin with IPTG-inducible expression |
